# Overcoming uncollapsed haplotypes in long-read assemblies of non-model organisms

**DOI:** 10.1101/2020.03.16.993428

**Authors:** Nadège Guiglielmoni, Antoine Houtain, Alessandro Derzelle, Karine van Doninck, Jean-François Flot

**Affiliations:** Service Evolution Biologique et Ecologie, Université libre de Bruxelles (ULB), 1050 Brussels, Belgium; Laboratoire d’Ecologie et Génétique Evolutive, Université de Namur, 5000 Namur, Belgium; Département de Biologie des Organismes, Université libre de Bruxelles (ULB), 1050 Brussels, Belgium; Interuniversity Institute of Bioinformatics in Brussels - (IB)^2^, 1050 Brussels, Belgium

**Keywords:** genome assembly, long reads, haplotypes collapsing

## Abstract

**Background:** Third-generation sequencing, also called long-read sequencing, is revolutionizing genome assembly: as PacBio and Nanopore technologies become more accessible in technicity and in cost, long-read assemblers flourish and are starting to deliver chromosome-level assemblies. However, these long reads are also error-prone, making the generation of a haploid reference out of a diploid genome a difficult enterprise. Although failure to properly collapse haplotypes results in fragmented and/or structurally incorrect assemblies and wreaks havoc on orthology inference pipelines, this serious issue is rarely acknowledged and dealt with in genomic projects, and an independent, comparative benchmark of the capacity of assemblers and post-processing tools to properly collapse or purge haplotypes is still lacking.

**Results:** To fill this gap, we tested different assembly strategies on the genome of the rotifer *Adineta vaga*, a non-model organism for which high coverages of both PacBio and Nanopore reads were available. The assemblers we tested (Canu, Flye, NextDenovo, Ra, Raven, Shasta and wtdbg2) exhibited strikingly different behaviors when dealing with highly heterozygous regions, resulting in variable amounts of uncollapsed haplotypes. Filtering out shorter reads generally improved haploid assemblies, and we also benchmarked three post-processing tools aimed at detecting and purging uncollapsed haplotypes in long-read assemblies: HaploMerger2, purge_haplotigs and purge_dups.

**Conclusions:** Testing these strategies separately and in combination revealed several approaches able to generate haploid assemblies with genome sizes, coverage distributions, and completeness close to expectations.

## Background

With the advent of third-generation sequencing, high-quality assemblies are now commonly achieved for all types of organisms. The rise of two main long-read sequencing companies, Pacific Biosciences (PacBio) and Oxford Nanopore Technologies (Nanopore), has prompted an increase in output as well as a decrease in cost, making their technologies more accessible to research teams and more applicable to any genome, including the more challenging ones. The primary advantage of long reads over short reads (such as those generated by Illumina sequencing platforms) is their typical length two or three orders of magnitude greater (Sedlazeck et al., 2018). As a result, long reads facilitate genome assembly into contigs and scaffolds as they can span repetitive regions (Pollard et al., 2018) and resolve haplotypes (Patterson et al., 2015).

However, long reads typically have a much higher error rate than Illumina data, and these errors are mainly insertions and deletions (vs. substitutions for Illumina reads). PacBio data have a random error pattern that can be compensated with high coverage, and recent developments have aimed to increase accuracy by generating circular consensus sequences (Wenger et al., 2019). Nanopore reads, on the other hand, have systematic errors in homopolymeric regions and therefore Nanopore contigs generally require further correction using Illumina and/or PacBio reads, in a process called “polishing” (Kundu et al., 2019; Zimin and Salzberg, 2020). Despite this disadvantage, Nanopore reads are currently much longer than PacBio reads, with runs attaining N50s over 100 kilobases (kb) and longest reads spanning over 1 Megabase (Mb) (Jain et al., 2018; Miga et al., 2019).

This progress has prompted the development of programs dedicated to producing *de novo* assemblies from long reads, all of which follow the Overlap Layout Consensus (OLC) paradigm (Miller et al., 2010). Briefly, OLC methods start by building an overlap graph (the “O” step), then simplify it and clean it by applying various heuristics (the “L” step), which typically include the removal of transitively inferable overlaps, and finally compute the consensus sequence of each contig (the “C” step). Some long-read assemblers follow strictly this paradigm, such as Flye (Kolmogorov et al., 2019), Ra (Vaser and Šikić, 2019), Raven (Vaser and Šikić, 2020), a further development of Ra by the same authors, Shasta (Shafin et al., 2020) and wtdbg2 (Ruan and Li, 2020); whereas other assemblers such as Canu (Koren et al., 2017) and NextDenovo (NextOmics, 2019) add a preliminary correction step based on an all-versus-all alignment of the reads (see Table 1). At present, most assemblers aim to generate a haploid assembly, in which each region of the genome is represented exactly once. For diploid or polyploid genomes, haploid assemblies include only one version of each heterozygous region and are therefore reduced representations of the actual complexity of the genome. Nevertheless, they provide a precious resource for genome analysis as they make it possible to compare easily genome structures, gene sets across species, identify orthologs for phylogenomic analysis, and to detect variants across individuals.

**Table 1.** Assemblers included in this study.

| Assembler | Reads correction | Version used | Access |
| --- | --- | --- | --- |
| Canu | ✓ | 1.9 | github.com/marbl/canu |
| Flye | × | 2.5 | github.com/fenderglass/Flye |
| NextDenovo | ✓ | 2.2 | github.com/Nextomics/NextDenovo |
| Ra | × | 0.2.1 | github.com/lbcb-sci/ra |
| Raven | × | 0.0.7 | github.com/lbcb-sci/raven |
| Shasta | × | 0.3.0 | github.com/chanzuckerberg/shasta |
| wtdbg2 | × | 2.5 | github.com/ruanjue/wtdbg2 |

Long-read assemblers were recently benchmarked on real and simulated PacBio and Nanopore bacterial datasets (Wick and Holt, 2019), and all assemblers tested proved their efficiency at reconstructing small haploid genomes within one hour and with a low RAM usage. For eukaryotic genomes, the wtdbg2 publication (Ruan and Li, 2020) included an evaluation of Canu, Flye, Ra and wtdbg2 on several model organisms (*Caenorhabditis elegans, Drosophila melanogaster, Arabidopsis thaliana* and *Homo sapiens*), all of which had low levels of heterozygosity (at most 1% for *A. thaliana*). This comparison suggested that wtdbg2 produces fewer artefactual duplications, but no attempt was made to pre-filter reads or post-process assemblies to improve the result. Hence, a comprehensive evaluation of strategies to generate a structurally correct haploid assembly of a non-model, heterozygous diploid organism is still lacking.

To fill this gap, we present here a quantitative and qualitative assessment of seven long-read assemblers on the relatively small eukaryotic genome of the bdelloid rotifer *Adineta vaga*, for which a draft assembly based on short reads was published some years ago (Flot et al., 2013). As with most non-model organisms, *Adineta vaga’s* genome presents a mid-range heterozygosity of ca. 2% with a mix of highly heterozygous and low-heterozygosity regions, making such genome more challenging to assemble than those of model organisms, which often exhibit very low levels of polymorphism (Leffler et al., 2012).

In addition to assessing the ability of these seven assemblers to collapse highly heterozygous regions, we investigated whether adding a pre-assembly read-filtering step (keeping only the longest reads) or removing uncollapsed haplotypes post-assembly (using existing tools HaploMerger2 (Huang et al., 2017), purge_dups (Guan et al., 2020) and/or purge_haplotigs (Roach et al., 2018)) improved the assembly. HaploMerger2 detects uncollapsed haplotypes in assemblies based on sequence similarity alone and can process both low and high-heterozygosity genomes. Along with sequence similarity, purge_dups and purge_haplotigs take into account the coverage depth obtained by mapping short and/or long reads to the contigs. Coverage depth represents the number of reads covering a position in a contig (computed after mapping reads on the assembly). The contigs are then aligned to select duplicates accurately and remove them. While purge_dups sets its coverage thresholds automatically, purge_haplotigs requires user-provided values. As the focus of the present benchmark was on dealing with uncollapsed haplotypes and not on polishing assemblies (a step for which many tools are available and that would represent a benchmark topic in itself), we did not perform polishing of our contigs.

Assemblies were evaluated using several metrics quantifying their level of continuity and the correctness of their haploid representation of a diploid genome: namely their assembly size, N50, BUSCO completeness, *k*-mer completeness, and coverage distribution. The assembly size represents the sum of the lengths of all the contigs in the assemblies. The N50 is a popular metrics that reflects directly the continuity of the assembly but does not account for possible structural errors; it is defined as the length of the largest contig for which 50% of the assembly size is contained in contigs of equal or greater length. The BUSCO completeness (Simão et al., 2015) assesses the number of orthologs retrieved completely from the assembly in one or several copies: a high-quality, properly collapsed haploid assembly should exhibit a high number of complete single-copy BUSCO features and a low number of duplicated BUSCO features. The *k*-mer completeness is the percentage of solid (i.e., frequently observed and therefore probably correct) *k*-mers in the set of reads present in the assembly. In the case of a haploid assembly of a diploid genome, all homozygous *k*-mers (i.e., *k*-mers that are shared by the two haplotypes) should be represented in the assembly, whereas only half of the heterozygous *k*-mers (i.e., *k*-mers that are found in only one haplotype) should be represented. To detect both underpurging and overpurging, we focused in our benchmark on the *k*-mer completeness of heterozygous *k*-mers: as we expect only half of them to be present in a haploid assembly, a well-collapsed assembly should exhibit a *k*-mer completeness of about 50%, whereas a lower value indicates that too many *k*-mers were lost (overpurging) and a higher value indicates that too many *k*-mers were retained (underpurging).

We also investigated the distribution of coverage depth of each assembly. In an ideal haploid assembly, all positions should be equally covered, hence we would expect a single peak in the coverage distribution. Based on our analysis of the coverage distribution, we developed a new metrics to evaluate the haploidy, or proper collapsing, of assemblies of diploid genomes. The haploidy score is based on the identification of two peaks in the per-base coverage depth distribution: a high-coverage peak that corresponds to bases in collapsed haplotypes (hereafter called “collapsed peak”), and a peak at about half-coverage of the latter that corresponds to bases in uncollapsed haplotypes (“uncollapsed peak”). The haploidy score represents the fraction of collapsed bases in the assembly, and is equal to C/(C+U/2), i.e. the ratio of the area of the collapsed peak (C) divided by the sum of the area of the collapsed peak (C) and half of the area of the uncollapsed peak (U/2). This metrics reaches its maximum of 1.0 when there is no uncollapsed peak, in a perfectly collapsed assembly, whereas it returns 0.0 when the assembly is not collapsed at all (as in the case of a phased diploid assembly). We implemented this metric in a new tool called HapPy (Houtain et al., 2020) available at github.com/AntoineHo/HapPy.

Finally, as computational resources can be a limiting factor in genome assembly, we compared the CPU time and RAM usage for the different assemblers tested by running them on the same machine under the same conditions. Canu and NextDenovo were not included in this comparison, as they required significantly higher resources and had to be run on different machines.

## Results

### Preliminary observations

The genome size of our benchmark organism was estimated as 102 Mb based on *k*-mer frequencies (see Methods). We initially ran each assembler five times on our complete and filtered Nanopore and PacBio datasets, but we observed that the N50 and BUSCO scores of the resulting assemblies were very similar to each other (Supplementary Figures S1-S2). As a result, we chose randomly one replicate assembly from each assembler and used this replicate for the subsequent haplotype-purging step.

To represent assembly statistics in a comprehensive manner, we defined four scores (see Methods): size, that is 1 minus the distance of the assembly size to the expected genome size; N50; completeness, a combined metrics that includes both the single-copy BUSCO score and the distance of the observed *k*-mer completeness to the expected value of 50%; haploidy, computed using HapPy.

### Assemblies of PacBio reads

Full assembly statistics are available in Figure S3 and Table S1-S2, whereas a summary of the results is presented in Figure 1.

**Figure 1.**
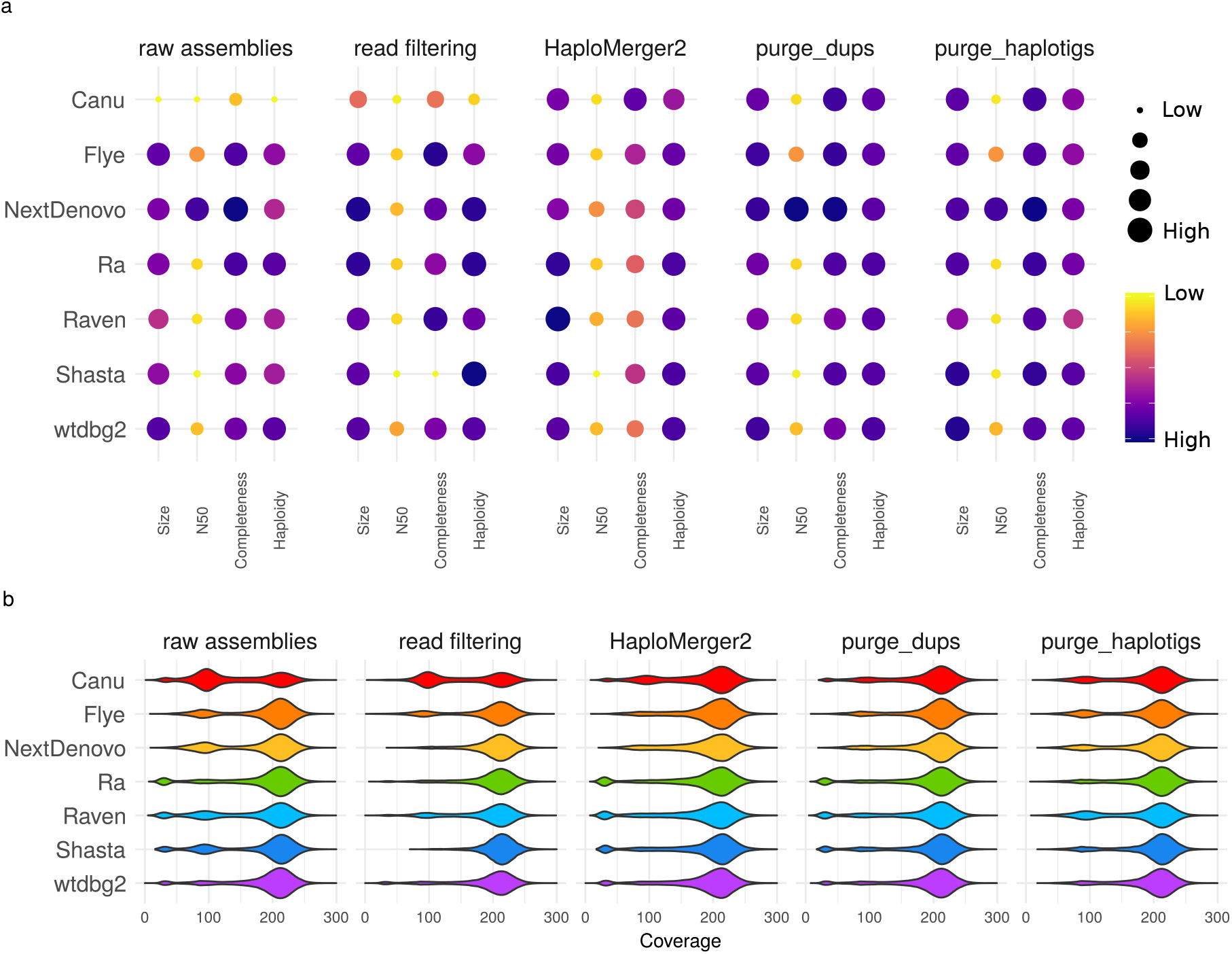
Statistics of raw assemblies obtained from the full PacBio dataset (raw assemblies), with a preliminary read-filtering step (keeping only reads larger than 15 kb), or a subsequent removal of uncollapsed haplotypes with HaploMerger2, purge_dups, or purge_haplotigs. a) Assembly scores for size, N50, completeness and haploidy. b) Long-read coverage distribution over the contigs.

#### Raw assemblies

All raw assemblies of the full PacBio datasets were oversized compared to the estimated genome size of 102 Mb, ranging from 114 Mb (wtdbg2) to 169 Mb (Canu) (Figure 1a, raw assemblies). The NextDenovo assembly obtained the highest N50 score with a value of 8.9 Mb, while other assemblies had an N50 ranging from 301 kb (Canu) to 2.7 Mb (Flye). The Canu assembly had the lowest completeness score, as its *k*-mer completeness was way above the expected value of 50% (73.54%) and its number of single-copy BUSCOs was only 538, compared to a highest value of 687 features (NextDenovo). The wtdbg2 and Ra assemblies scored the highest according to the haploidy metrics (0.90), while Canu obtained the lowest score (0.59).

Larger assembly sizes correlated with highly bimodal coverage distributions (Figure 1b, raw assemblies). This was particularly the case with Canu assemblies, which exhibited two large peaks plus a smaller, low-coverage peak. The high-coverage peak, around 210X, was the collapsed peak C whereas the 100X peak, at about half-coverage of the C peak, corresponded to the uncollapsed peak U. In the case of the Canu assembly, the U peak was larger than the C peak, revealing a poor collapsing of highly heterozygous regions. The Flye, NextDenovo, Raven and Shasta assemblies also exhibited two peaks in their coverage distribution, although their U peak was smaller than the one of Canu. The Ra, Raven, Shasta, and wtdbg2 assemblies exhibited an additional low-coverage peak identified as contaminants (see Supplementary Figures S4-S10).

#### Read filtering

Selecting a subset of the longest PacBio reads (over 15 kb) resulted in assemblies closer to the expected size than assemblies of all reads for Canu (145 Mb), NextDenovo (99 Mb), Ra (108 Mb), and Raven (117 Mb) (Figure 1a, read filtering). In the case of NextDenovo, read filtering made the N50 assembly drop to a value comparable with other assemblies (from 8.9 to 1.8 Mb). Most assemblies maintained their completeness score, and it even increased for some (Canu, Flye, Raven). The Shasta assembly was shorter than the expected size (89 Mb), possibly due to the sub-optimal sequencing depth of the filtered read set (as Shasta is tuned for a 60X-depth) and its completeness score was the lowest among all assemblies. As for the coverage distribution (Figure 1b, read filtering), the Ra assembly no longer showed a low-coverage contaminant peak, and for the NextDenovo assembly the U peak was absent. The Raven assembly also displayed a strong C peak, but the U peak remained, although reduced.

#### Haplotig purging

When adding a post-assembly haplotig-purging step, we observed strikingly different results depending on the combination of assembler and post-assembly purging tool, namely HaploMerger2, purge_dups, purge_haplotigs. While HaploMerger2 reduced the size of all assemblies (resulting in higher size scores on Figure 1a), it also led to a decrease of the completeness score of all assemblies, except for Canu (Figure 1a, HaploMerger2). Nevertheless, the haploidy scores all increased (with a minimum of 0.84 for Canu and a maximum of 0.92 for Ra and wtdbg2) as U peaks decreased drastically in all coverage distributions (Figure 1b, HaploMerger2). Assemblies purged with purge_dups were all closer to the expected size of 102 Mb, and the N50 and completeness scores were maintained or even improved (Figure 1a, purge_dups). The haploidy scores were also improved as they ranged from 0.89 (Canu and Flye) to 0.91 (Ra and wtdbg2). The coverage distributions showed that the U peaks were removed or at least reduced, but the low-coverage peaks were not (Figure 1b, purge_dups). After purging with purge_haplotigs, all assembly sizes were closer to the expected size except for Flye, and the N50 and completeness scores were maintained or even improved (Figure 1a, purge_haplotigs). The haploidy scores were improved for Canu, NextDenovo and Shasta. The coverage distributions showed a reduction of the U peak for the Canu, NextDenovo and Shasta assemblies, explaining their higher haploidy scores (Figure 1b, purge_haplotigs). The low-coverage peaks were removed for Ra, Raven, Shasta and wtdbg2, explaining their smaller assembly size.

#### Combination of read filtering and haplotig purging

The combination of read filtering and haplotig purging resulted in assemblies that almost all had a unimodal coverage distribution (Figure S11). As observed with assemblies of all reads, HaploMerger2 seemed to overpurge assemblies of the filtered datasets. Only the Canu assembly remained above the expected size (108 Mb), but its statistics were similar to those obtained with the Canu assembly of all reads. Assemblies purged with purge_dups or purge_haplotigs were closer to the estimated size and exhibited high numbers of single-copy BUSCO features, although the Shasta assembly of filtered reads was already too small prior to purging (89 Mb). The combination of pre-assembly read filtering and post-assembly purge_dups seemed beneficial for most assemblers with exception of Shasta, as the resulting assemblies ranged in haploidy from 0.90 (Canu and Flye) to 0.97 (NextDenovo). By contrast, the improvements observed when using a combination of read filtering and purge_haplotigs were similar to those obtained with either one of the two. NextDenovo, Ra and wtdbg2 assemblies had satisfying assembly sizes (from 99 to 107 Mb), high haploidy scores (from 0.85 to 0.96) and unimodal coverage distributions, but similar scores were also obtained using only read selection for NextDenovo and Ra and using only purge_haplotigs for wtdbg2.

#### Combination of haplotig-purging tools

The combination of purge_dups or purge_haplotigs with HaploMerger2 resulted in problems similar to those observed on assemblies purged only with HaploMerger2: except for Canu, assemblies were shorter than expected and/or the number of single-copy BUSCO features dropped. However, the Canu assembly purged with both HaploMerger2 and purge_dups had a high number of single-copy BUSCOs (672 features), a *k*-mer completeness close to 50% (51.73%), and no half-coverage peak (Figure S12). Assemblies purged with both purge_dups and purge_haplotigs were all reduced in size, but none went below the expected size. The BUSCO score and the *k*-mer completeness were stable, and the haploidy scores ranged from 0.88 (Canu and Raven) to 0.92 (NextDenovo). The coverage distributions were all close to a unimodal distribution, as there were no low-coverage contigs and the U peaks had mostly disappeared.

### Assemblies of Nanopore reads

Full assembly statistics are provided in Figure S13 and Table S3-S4.

#### Raw assemblies

Similarly to the PacBio assemblies, Nanopore assembly sizes exceeded the expected size of 102 Mb, ranging from 118 Mb (Shasta, wtdbg2) to 154 Mb (Canu) (Figure 2a, raw assemblies). Nanopore assemblies achieved a much higher continuity than PacBio assemblies, as PacBio assemblies had a lowest N50 of 301 kb while Nanopore assemblies had a lowest N50 1.6 Mb. Canu, NextDenovo, Ra and wtdbg2 achieved a N50 over 5 Mb using Nanopore reads. The number of complete single-copy BUSCOs was lower in Nanopore assemblies (up to 559) than in PacBio assemblies (up to 699). The *k*-mer completeness was also usually lower for Nanopore assemblies, ranging from 38.61 to 54.82%, compared to 47.71-73.54% for PacBio assemblies. These lower values for *k*-mer completeness in Nanopore assemblies were likely not due to a better collapsing but rather to systematic errors. The haploidy scores were higher for Nanopore assemblies produced by Canu, Ra, Raven, Shasta and wtdbg2, in comparison with PacBio assemblies, but this score was lower for Flye and NextDenovo assemblies (Table S1,S3). The coverage distribution of the Canu assembly exhibited two distinct U (uncollapsed) and C (collapsed) peaks respectively around 75X and 160X, indicating that many haplotypes were not collapsed and were therefore represented twice in the assemblies (Figure 2b, raw assemblies). This was also the case for the Flye, NextDenovo, Raven and Shasta assemblies, albeit their U peak was smaller than the one of the Canu assembly. The Ra, Raven, Shasta and wtdbg2 assemblies had an additional low-coverage peak identified as contaminations (see Supplementary Figures S14-S20).

**Figure 2.**
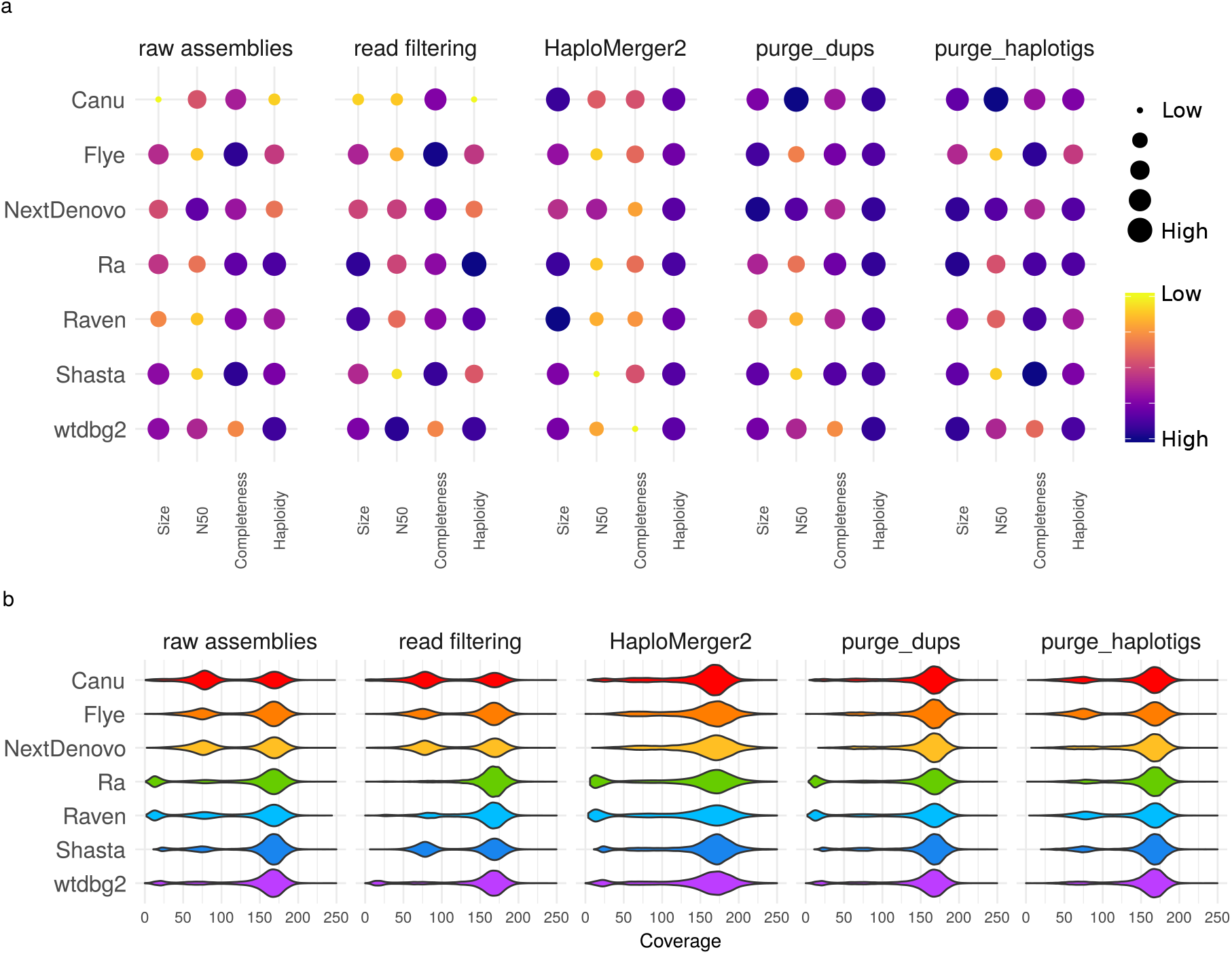
Statistics of raw assemblies obtained from the full Nanopore dataset (raw assemblies), with a preliminary read filtering step (keeping only reads larger than 30 kb) or a subsequent removal of uncollapsed haplotypes with HaploMerger2, purge_dups, or purge_haplotigs. a) Assembly scores for size, N50, completeness and haploidy. b) Long-reads coverage distribution over the contigs.

#### Read filtering

When assembling a subset of the longest Nanopore reads (over 30 kb), the Ra and Raven assemblies had sizes closer to the estimated size (97 and 109 Mb) and did not exhibit a contaminant peak in their coverage distribution, whereas other assemblies were unmodified (Figure 2, read filtering). The Raven assembly of filtered reads had a smaller U peak compared to the raw assembly, but it was still present, whereas the U peak of the Ra assemby disappeared. These improvements came along with increased haploidy scores: from 0.90 to 0.95 for Ra, and from 0.83 to 0.89 for Raven. The result produced by Shasta with read filtering was the opposite, as the size and haploidy scores became lower and the U peak became larger. This increase in size of the U peak explained the higher *k*-mer completeness obtained due to a higher percentage of uncollapsed homozygous and heterozygous *k*-mers (Figure S21-S22). Overall, read filtering did not significantly affect completeness scores.

#### Haplotig purging

As we observed with PacBio reads, all assembly sizes were reduced by HaploMerger2, which resulted in higher size scores, and the U peaks were removed from the coverage distributions, but this also led to lower scores for completeness (Figure 2, HaploMerger2). purge_dups improved size scores while maintaining or increasing completeness scores and removing U peaks (Figure 2, purge_dups). These improved coverage distributions resulted in higher haploidy scores, ranging from 0.90 (Flye and Raven) to 0.93 (Ra and Raven). purge_haplotigs improved size scores for all assemblies except Flye and also kept high completeness scores (Figure 2). Haploidy scores were higher for assemblies produced by Canu and NextDenovo, and the coverage distribution shows that U peak were indeed reduced or removed for Canu and NextDenovo assemblies, while the contaminant peaks were removed for Ra, Raven, Shasta and wtdbg2 assemblies. We observed again that HaploMerger2 generally decreased the continuity and quality metrics of the assemblies, while purge_dups and purge_haplotigs did not. Interestingly, the Canu assembly obtained the highest N50 of all the assemblies presented in this paper (12.4 Mb) when raw assemblies were purged with either purge_dups or purge_haplotigs.

#### Combination of read filtering and haplotig purging

The read-filtered (> 30kb) Nanopore assemblies after HaploMerger2 were too short (72-87 Mb), except for the Canu assembly (107 Mb; Figure S23). All assemblies of the longest reads after purge_dups were close to the expected genome size (96-109 Mb), except for the wtdbg2 assembly (114 Mb), and the coverage distributions were almost all unimodal, with a haploidy score ranging from 0.87 (Canu) to 0.96 (Ra). Notably, the Shasta assembly of filtered reads had a strong U peak that was completely removed by purge_dups, which brought the assembly to a size close to our estimation (102 Mb) and resulted in a high number of singe-copy BUSCO features (519 features). purge_haplotigs was apparently less efficient, as the U peak was reduced for the Canu and Raven assemblies but remained high for the Flye, NextDenovo and Shasta assemblies. The BUSCO scores of assemblies purged with purge_dups or purge_haplotigs were similarly high (up to 539 single-copy BUSCO features, Ra).

#### Combination of haplotig-purging tools

The combination of HaploMerger2 with either purge_dups or purge_haplotigs led to excessively shorter assemblies, except for Canu, as HaploMerger2 tended to overpurge (Figure S24). The assemblies purged with both purge_dups and purge_haplotigs were all close to the expected size (100-110 Mb), their single-copy BUSCO score was maintained (up to 559 features, Flye assembly), they had a *k*-mer completeness below 50%, their haploidy ranged from 0.90 (Flye and Raven) to 0.94 (NextDenovo) and their coverage distribution was unimodal or close to it.

### Impact of sequencing depth

We further evaluated the impact of sequencing depths ranging from 10X to 100X on the size, N50, BUSCO score, and haploidy metrics of PacBio and Nanopore assemblies (Figure 3).

**Figure 3.**
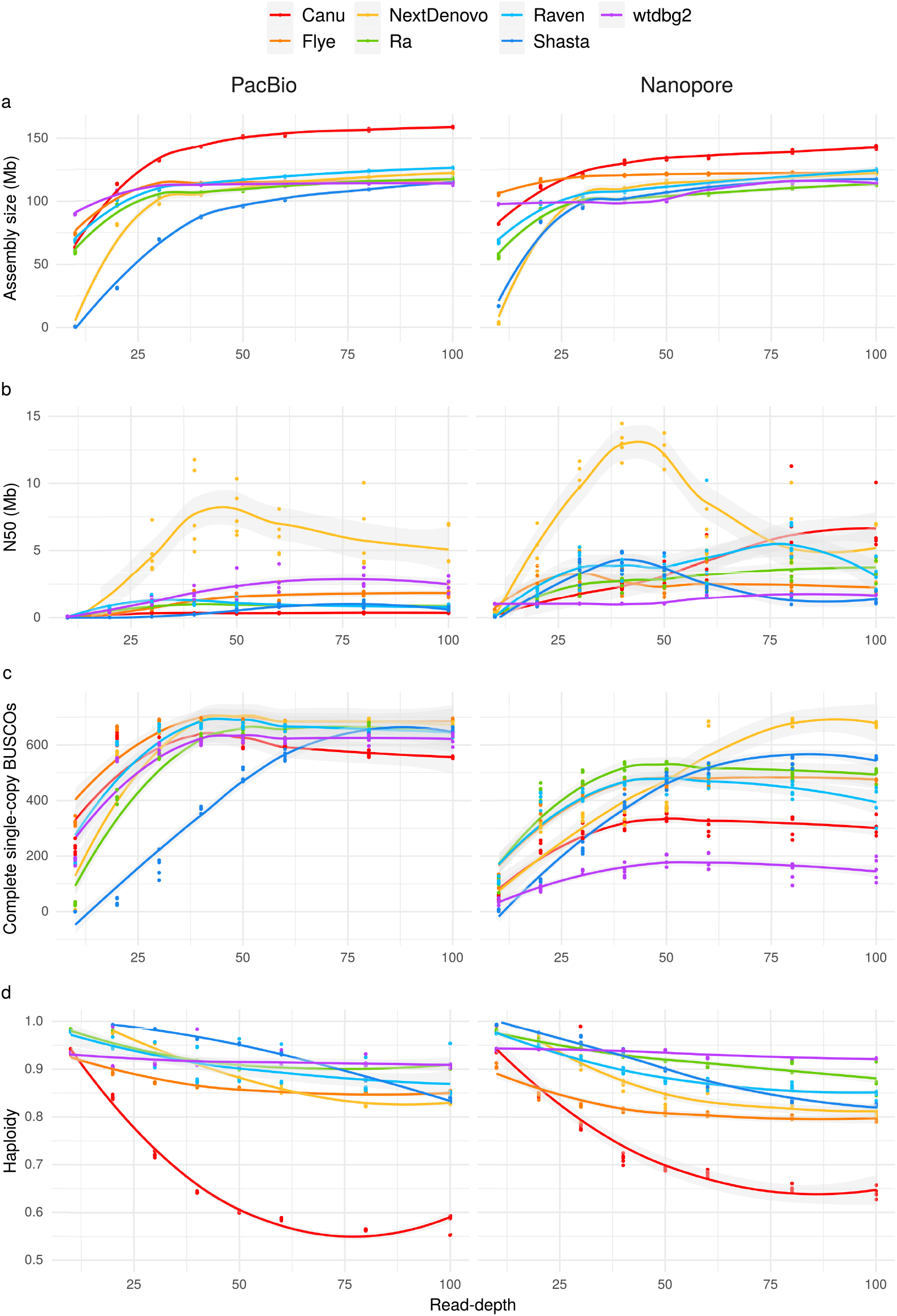
Statistics of the PacBio and Nanopore assemblies depending on sequencing depth, with a) assembly size, b) N50, c) complete single-copy BUSCOs and d) haploidy. The assemblies were ran on five random subsamplings of the long-read datasets.

Almost no assembler produced decent outputs with a sequencing depth of 10X, under which condition NextDenovo and Shasta performed worst. However, with wtbdg2 on PacBio and Nanopore reads, and with Flye on PacBio reads, the assembly size was close to the expected size. At 20X, Canu, Flye, Ra, Raven and wtdbg2 reached at least 93 Mb. Assembly sizes increase sharply up to 40X, except for Canu for which the assemblies kept increasing in size with sequencing depth.

While there was no variation in assembly size among replicates, N50s were highly variable for PacBio assemblies with NextDenovo, Flye and wtdbg2, and for almost all Nanopore assemblies other than wtdbg2. For NextDenovo, there was a clear optimal coverage at about 40X in terms of N50 with PacBio as well as Nanopore reads. Other assemblers also showed a decrease in N50 above a certain sequencing depth: for PacBio assemblies, this happened for Raven over 30X and for Shasta and wtbdg2 over 80X; for Nanopore assemblies, this happened for Flye over 30X, for Shasta over 40X, and for Raven over 80X.

The number of single-copy BUSCO features did not vary much among replicate subsamplings; only NextDenovo exhibited some variability in that regard. Over 40X, the number of single-copy BUSCOs became strikingly stable for most assemblers with both PacBio and Nanopore reads, except for Shasta that is tuned for a 60X depth. For Canu assemblies, the number of complete single-copy BUSCOs decreased when sequencing depth exceeded 30X. For wtdbg2 the complete single-copy BUSCOs remained low for the Nanopore dataset at the different sequencing depths.

For most assemblers, haploidy decreased when sequencing depth increased, and this was especially drastic for Canu assemblies. Only the assemblies produced by wtdbg2 had a stable haploidy value, whereas the Ra assemblies of PacBio reads also exhibited limited variations in haploidy.

### Computational performance

We evaluated the computational performance of Flye, Ra, Raven, Shasta and wtdbg2. Canu and NextDenovo were not included in this comparison as they required higher resources and were therefore run on different machines. RAM usage and/or CPU time (Figure 4) increased with sequencing depth, except for wtdbg2 that did not display much change. wtdbg2 was the assembler that required the lowest amount of RAM (less than 20 GB) and was the second fastest on PacBio reads (less than 53 hours) and Nanopore reads (less than 192 hours). Flye was the least efficient among the assemblers that could be included in this comparison. Its RAM usage was the highest for Nanopore assemblies, with high variability. It also had the longest CPU time for both PacBio and Nanopore reads. Notably, Shasta generally required a lot of RAM and had the highest memory footprint for high-coverage PacBio data, but it ran the fastest on PacBio as well as on Nanopore reads.

**Figure 4.**
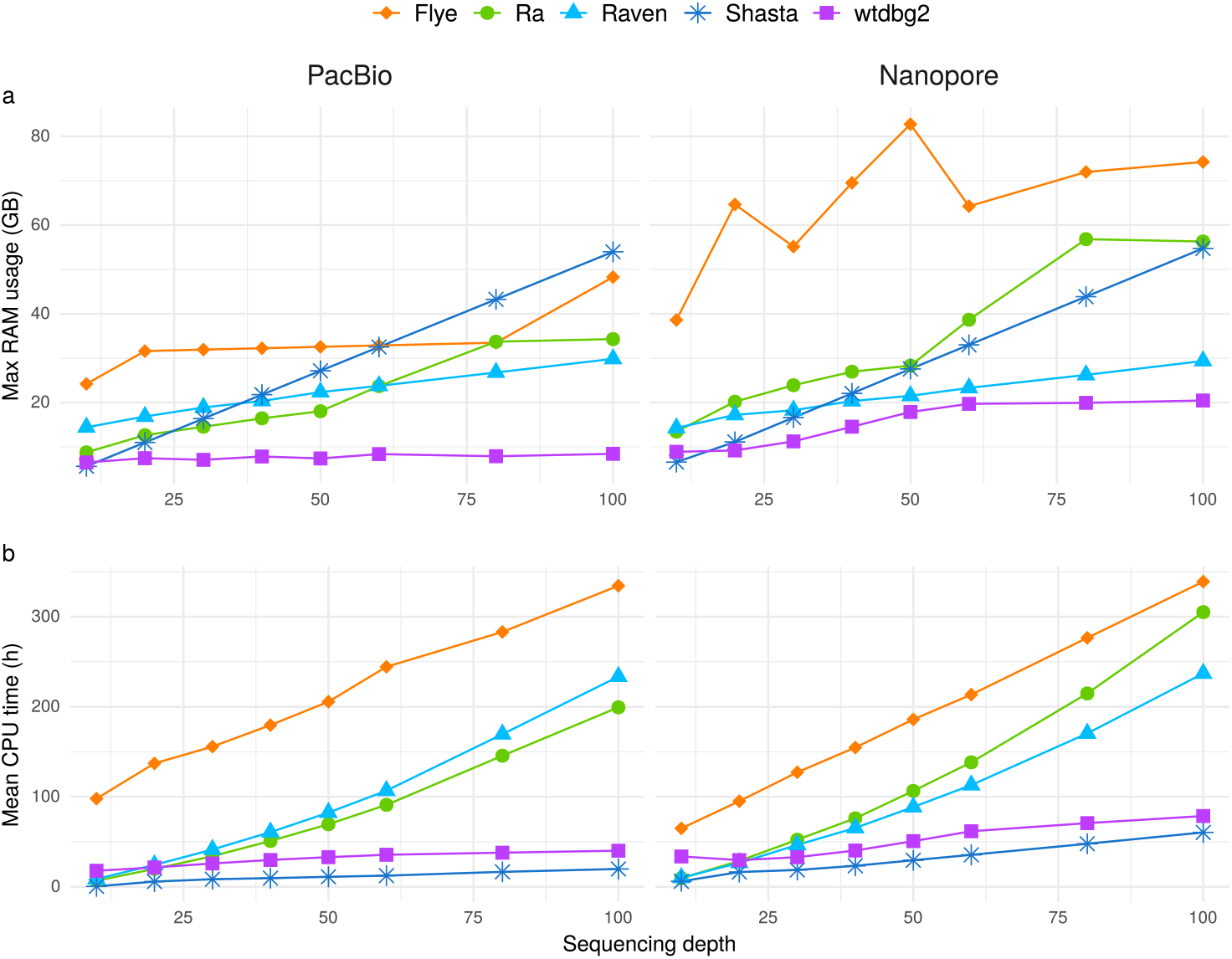
Computational resources (RAM and CPU time) used by the assemblers. a) Maximum RAM usage and b) mean CPU time depending on sequencing depth. Canu and NextDenovo were not included in this comparison as they were run on different machines.

## Discussion

### Comparison of PacBio and Nanopore assemblies

While PacBio assemblies were superior in terms of completeness, the continuity of Nanopore assemblies was far greater for most assemblers, probably due to the greater length of Nanopore reads. The lower completeness scores of Nanopore assemblies likely resulted from the lower accuracy along with the non-random error pattern of Nanopore reads producing errors (mostly indels) in the consensus sequences produced by the assemblers. These systematic errors in Nanopore reads may be improved with more recent protocols (Van der Verren et al., 2020) and basecallers (Wick et al., 2019).

However, there was no striking difference regarding the efficiency of haplotype collapsing when assembling PacBio or Nanopore reads. The results in terms of coverage distribution and haploidy were similar, and it appears therefore that both technologies can be used to produce properly collapsed assemblies. An important finding was that keeping only reads longer than a certain threshold improved in many cases the quality of haploid assemblies, as it led to a decrease in coverage depth and to a lower support for both haplotypes, thereby favoring the collapse of the region. We further observed that read filtering did not lead to a decrease in assembly quality as long as the sequencing depth remained sufficient, and for some assemblers read filtering resulted in an increase in N50 and/or in completeness. Reads may also be selected using both length and quality as criteria using the tool Filtlong (Wick, 2017), which can additionally use short reads to help trim low-quality regions; however, this approach was not tested in our benchmark.

### Successful combinations for haploid assemblies

We found that Canu poorly collapsed alleles and yielded oversized assemblies. The program did not seem able to collapse highly divergent regions on its own. Post-processing assemblies using haplotype-purging tools greatly improved haploidy. All three tools tested reduced the assembly size and the size of the U peak, but purging was most efficient using purge_dups. Purging Canu assemblies with purge_dups or purge_haplotigs improved greatly their haploidy. Besides, Canu assemblies of Nanopore reads purged with purge_dups or purge_haplotigs yielded the highest N50s among all our tests. However, the computational resources required to run Canu may be a limiting factor: although the RAM usage can be limited with the parameter maxMemory, this reduced the number of CPUs used and increased the running time.

Flye assemblies exhibited uncollapsed haplotypes too; selecting the longest reads did not help, but HaploMerger2 and purge_dups improved collapsing. While Flye assemblies purged with HaploMerger2 were shorter than the expected size and suffered from a decrease in completeness, those purged with purge_dups had an assembly size close to the expectation and kept a good quality. Thus, Flye and purge_dups appeared as a good combination for generating a haploid assembly.

NextDenovo produced the assemblies with the highest N50s before post-processing, but with poorly collapsed haplotypes. This problem was alleviated for PacBio assemblies when selecting the longest reads, and uncollapsed haplotypes were efficiently removed by haplotig-purging tools. The best haploidy values were achieved when combining NextDenovo with read filtering, purge_dups and/or purge_haplotigs. These assemblies also reached high values of continuity and completeness. However, although NextDenovo runs quickly, it requires a large amount of RAM.

Ra was more efficient at collapsing haplotypes than most assemblers, and its oversized assemblies were rather due to contaminants. Ra assemblies were therefore not much improved by the post-processing haplotype merging tools. Ra assemblies proved even better when using only the longest reads, which led to better continuity, equal completeness and improved collapsing. Although Ra was not the most computationally efficient assembler, its RAM usage and CPU time remained low up to a sequencing depth of 50X-depth; thus it appeared even more desirable to use only a subset of the longest reads for this assembler.

Although Raven is a further development of Ra, it exhibited a different behavior: Raven was more computationally efficient but produced less collapsed haplotypes compared to Ra. Read filtering and/or haplotig-purging reduced these uncollapsed haplotypes, resulting in high-quality assemblies when combining read filtering with purge_dups, or with purge_dups and purge_haplotigs.

We observed singular results with Shasta. The assembly of filtered PacBio reads obtained with this tool were well collapsed but incomplete, likely due to an insufficient sequencing depth. On the other hand, the assembly of filtered Nanopore reads was less collapsed than the assembly of all reads. purge_dups efficiently purged the assemblies of all PacBio and Nanopore reads but, surprisingly, the best haploid Shasta assembly was obtained from filtered Nanopore reads purged with purge_dups. Shasta assemblies generally achieved a good completeness, but their continuity was lower than with other programs, as the developers explicitly aimed for quality over continuity.

wtdbg2 performed well on PacBio data, but less on Nanopore reads, for which it obtained the lowest completeness scores. This program did not seem to have difficulties with heterozygous regions, but low-coverage contigs identified as contaminants remained in the final assemblies. Read selection on size did not significantly improve the assemblies, but purge_haplotigs removed contaminant contigs, therefore improving the output. Short-read polishing would certainly improve the low completeness of Nanopore assemblies. Users may want to test this assembler as it collapses genomes well and runs fast using a moderate amount of RAM.

Based on the above, we recommend users interested in generating the best haploid assembly of a diploid genome to try all or some of the solutions described in Table 2, depending on the size of the genome they want to assemble, the technology of reads they have, and their available computational resources.

**Table 2.**
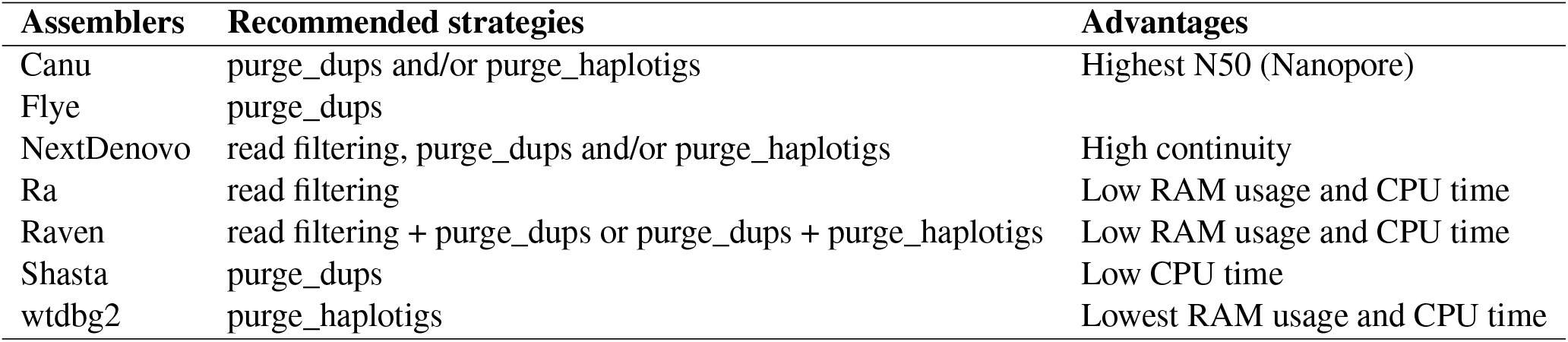
Recommended strategies for generating high-quality haploid assemblies.

### Impact of sequencing depth

Our study of the impact of sequencing depth on the assemblies showed that deeper sequencing usually did not result in higher continuity or improved haploidy of the assemblies. Most programs reached the expected assembly size between 10X and 30X, while the BUSCO scores and *k*-mer completeness plateaued around 40X. Depending on the assembler, N50s also decreased when sequencing depth went beyond a specific threshold. A deeper coverage may lead to erroneous low-coverage contigs and/or provide more support to both haplotypes in highly heterozygous regions, promoting incomplete collapsing of haplotypes. We observed in our benchmark that a deeper sequencing lead to a lower haploidy (as computed using HapPy). The combination of the continuity and quality metrics show that a sequencing depth of 40X is sufficient for generating a high-quality haploid assembly. Besides, a counter-intuitive finding is that a larger amount of reads does not improve the assembly and can even make it worse in terms of continuity and haploidy. Most assemblers seem optimized for sequencing depths around 30 to 40X and therefore did not appear to benefit from more data, except Shasta that is optimized for 60X.

### Assembly evaluation

We propose here a set of metrics to evaluate thoroughly genome assemblies and identify uncollapsed haplotypes. The N50, BUSCO score and *k*-mer completeness are commonly used to estimate the continuity and completeness of assemblies, but do not discriminate properly collapsed haploid assemblies. It is possible to combine the completeness and continuity with a comparison of assembly size vs. expected genome size and an examination of the coverage distribution to identify the best assemblies. We further described a new metric of the haploidy of an assembly and implemented it in HapPy. We used the haploidy metric to systematically evaluate haploid assemblies. HapPy gives an accurate numerical representation of the coverage distribution.

These metrics have their limits, and not one of them is sufficient to identify the best assembly. The N50 is the most popular metric to describe contigs as it represents the continuity, yet high N50s can be achieved with efficient scaffolding methods. Therefore, we should aim for high-quality contigs that can be later turned into high-quality scaffolds. Comparing the assembly size to an estimated genome size depends on the reliability of the estimation itself. Genome size can be estimated computationally with a *k*-mer spectrum (Mapleson et al., 2016; Ranallo-Benavidez et al., 2020), or experimentally using flow cytometry and Feulgen densitometry (Mulligan et al., 2014). The BUSCO score only represents orthologs and does not account for the completeness of non-coding regions. Besides, the *k*-mer completeness is not sufficient as a value, as it could reach 50% with an assembly that has a balanced combination of missing regions and uncollapsed ones. To better estimate the *k*-mer completeness, it is necessary to examine plots provided by KAT. Collapsed repeated regions appear in the coverage plot along the contigs as localized peaks of elevated coverage. Since most assemblers included in our benchmark can produce an assembly graph, we strongly advise readers to investigate this file using dedicated tools (Wick et al., 2015): uncollapsed haplotypes are usually observed as bubbles in the graph. Ideally, the contigs should be evaluated with all metrics available to find, if not the perfect assembly, the best assembly, while assessing its limitations.

Genome papers present only the most successful assembly strategy, but researchers usually try more than one method to obtain the best result. They rarely report details of all the tested approaches in publications, although such negative results could help the community and could guide developers in how to improve their tools. To improve on this situation, we suggest reporting alternative assembly results as supplementary material or separately on a preprint platform.

### Comparison with other studies

Few comparisons of long-read assemblers are currently available. A thorough study was conducted on simulated and real bacterial datasets (Wick and Holt, 2019), and showed that all these assemblers can achieve assemblies of varying quality. Flye and Raven were also tested in this study and emerged among the most reliable assemblers. We also found that these assemblers reached high single-copy BUSCO scores, but when processing data from a diploid organism, they are not the most efficient at collapsing haplotypes. Besides, the benchmark on bacterial datasets also showed that Canu required more computational time and memory than most assemblers. Although this benchmark gave essential information on the performance of these assemblers, eukaryotic genomes represent a completely different challenge. Recently, a publication compared different long-read sequencing technologies to assemble a plant genome, *Macadamia jansenii* (Murigneux et al., 2020). They included statistics for different assemblers and obtained, depending on the tool, oversized assemblies combined with heightened numbers of duplicated BUSCO features on an 80X PacBio dataset, while they did not observe such differences on a 30X Nanopore dataset. These results agree with our observations on the impact of sequencing depth, as the 30X Nanopore dataset was not problematic, while the 80X PacBio dataset was, likely because of a deeper sequencing.

### Toward high-quality diploid and polyploid assemblies

Haploid assemblies of multiploid (i.e. diploid or polyploid) organisms provide a partial representation of their genomes as only one version of all heterozygous regions is included in the assembly. Ideally, we would prefer to generate phased multiploid assemblies. Low-accuracy long reads can separate haplotypes for highly heterozygous regions, but their high error rates do not allow the identification and separation of small heterozygous regions. Furthermore, phased assemblies bring an extra challenge, as alleles from different heterozygous regions need to be correctly associated. A protocol for high-accuracy long reads (above 99%) has been released recently, called PacBio HiFi (Wenger et al., 2019), and brings new possibilities for phased multiploid assemblies. To better accommodate these high-accuracy long reads, new versions of assemblers have been released such as HiCanu (Nurk et al., 2020) (a development of Canu), hifiasm (Cheng et al., 2020), or Flye’s new option `--pacbio-hifi`. Fully phased assemblies will provide complete representations of multiploid genomes.

## Conclusion

We believe that benchmarks such as ours are essential to help researchers working on non-model organisms select a long-read sequencing technology and an assembly method suitable for their project. It will also help them understand better the results they obtain and in general improve the rapidly evolving field of genomics.

## Methods

All command lines are provided in Supplementary Table S5.

### Genome size estimation

The genome size of *Adineta vaga* was estimated using KAT (Mapleson et al., 2016) on an Illumina dataset of 25 millions paired-end 250 basepairs (bp) reads (see Table S2). The diploid size was estimated to 204.6 Mb, therefore a haploid assembly should have a length around 102.3 Mb.

### Long-read assemblies

Canu, Flye, NextDenovo, Ra, Raven, Shasta and wtdbg2 were tested on two *Adineta vaga* long-read datasets: PacBio reads totalling 23.5 Gb with a N50 of 11.6 kb; and Nanopore reads totalling 17.5 Gb with a N50 of 18.8 kb (after trimming using Porechop, github.com/rrwick/Porechop). All assemblers were used with default parameters, except for Shasta for which the minimum read length was set to zero (instead of the default 10 kb setting). To run Shasta on PacBio reads, we used the recommended parameters `--Assembly.consensusCaller Modal --Kmers.k 12`. When assemblers required an estimated size, the value 100 Mb was provided. PacBio assemblies were run on all reads and on reads > 15 kb (4.7 Gb). Nanopore assemblies were run on all reads and on reads > 30 kb (5.7 Gb). For both datasets, we tested several length thresholds to find the optimal one. For more details on the long-reads datasets we used, see the publication by Simion et *al*. (Simion et al., 2020) and Supplementary Table S6. To test for reproducibility, all assemblers were run five times.

### Purging duplicated regions

Reads were mapped on assemblies using minimap2 (Li, 2018). For each assembly, we ran `purge_haplotigs hist` (Roach et al., 2018) to compute coverage histograms that we used to set low, mid and high cutoffs; these values were then used by `purge_haplotigs cov` to detect suspect contigs. Finally, we ran `purge_haplotigs purge` to eliminate duplicated regions.

purge_dups (Guan et al., 2020) was run following instructions by first generating the configuration file and then purging the assembly. HaploMerger2 (Huang et al., 2017) was run by sequentially running the modules BuildDatabase, RepeatModeler, RepeatMasker and finally the main script of HaploMerger2 to purge the assembly.

### Impact of sequencing depth

To find out the impact of sequencing depth, five replicate subsets were randomly sampled from the long-read datasets using the script reformat.sh from BBTools, available at sourceforge.net/projects/bbmap/, by providing the desired number of bases. The assemblers were run on these subsets with the same parameters as previously, with the exception of Canu: the parameters stopOnLowCoverage was set to 1 to allow runs on low-depth datasets. We tested subsets with a sequencing depth of 10X, 20X, 30X, 40X, 50X, 60X, 80X, 100X.

### Assembly evaluation

To evaluate the assemblies, we ran BUSCO 4 (Simão et al., 2015) against metazoa odb10 (954 features) without the parameter `--long`. We ran `KAT comp` (Mapleson et al., 2016) to calculate *k*-mer completeness by reference to the same Illumina 2*250 bp dataset used to estimate the genome size. To compute coverage, long reads were mapped on one replicate assembly per assembler using minimap2 and the coverage was computed with tinycov, available at github.com/cmdoret/tinycov, with a window size of 20 kb.

### Haploidy evaluation

To evaluate the collapsing of assemblies based on the coverage distribution, we developed a script, available at github.com/AntoineHo/HapPy). HapPy estimates the haploidy of an assembly (a measure of how well it is collapsed) by analyzing the per-base coverage histogram obtained after mapping reads to the assembly. For a well-collapsed haploid assembly, this histogram should consists of one peak around the theoretical average depth of coverage.

Actual coverage curves rarely fit the theoretical expectations because of several factors:

– contaminant contigs (e.g. bacteria and viruses that were sequenced along the organism of interest) can appear as other peaks at an unexpected coverage, usually lower than the actual sequencing depth;
– some contigs or contig regions in the assembly may actually correspond to uncollapsed haplotigs;
– large and/or abundant hemizygous deletions, as well as haploid chromosomes (e.g. the Y chromosome of male mammals), can result in a half-coverage peaks; in such case, this peak is a biological signal that will prevent reaching a perfect haploidy score.

`HapPy` expects up to three peaks in the per-base coverage histogram: a low-coverage contaminant peak (if any), an uncollapsed peak at around half of the expected coverage and a collapsed peak around the expected coverage. `HapPy` first applies a smoothing on the curve with a Savitzky–Golay filter (Savitzky and Golay, 1964) then measures peaks height and widths using SciPy. Finally, `HapPy` computes the areas of the relevant peaks.

We defined an haploidy score using the following Equation 1, in which *U* is the area of the uncollapsed peak and C the area of the collapsed peak reported by `HapPy`.

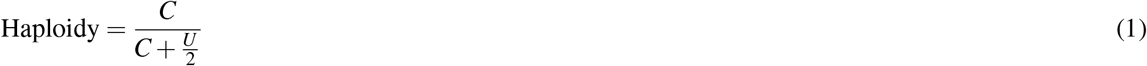

### Scoring

We defined four scores to evaluate the assemblies in Figures 1 and 2: size, N50, completeness and haploidy. The N50 score corresponds to the regular N50 value, and the haploidy score is computed using HapPy. The size score reflects the distance of the assembly size to the estimated haploid genome size and is computed following Equation 2, in which s is the assembly size and *G* the estimated haploid genome size (102 Mb).

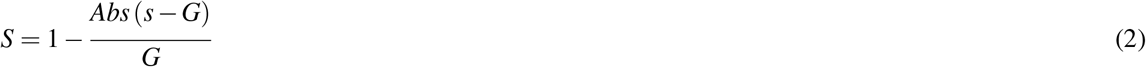

The completeness score includes both the number of single-copy BUSCO features and a measure of the distance of the observed *k*-mer completeness compared to the expected one. This metrics is computed using Equation 3, in which *k_obs_* is the observed *k*-mer completeness, *k_exp_* the expected *k*-mer completeness and *k_exp_* = 50.

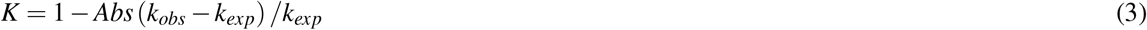

The number of single-copy BUSCOs and the value *K* are normalized on a 0 to 1 scale following Equation 4, in which *x_i_* is the initial value, *x_min_* the minimum value for all PacBio or Nanopore assemblies, and *x_max_* the maximum value for all PacBio or Nanopore assemblies.

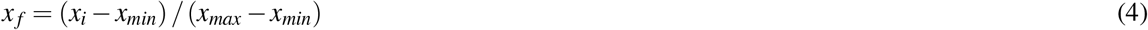

The final completeness score is computed using Equation 5, in which *B_norm_* is the normalized single-copy BUSCO score and *K_norm_* is the normalized *k*-mer completeness value computed previously.

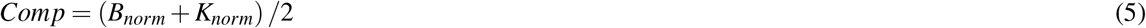

### Performance evaluation

For Flye, Ra, Raven, Shasta and wtdbg2, maximal RAM usage and mean CPU time were measured using the command time with 14 threads on a computer with an i9-9900X 3.5 Ghz processor and 128 GB RAM. NextDenovo was run on a computer with an Intel Xeon E5-2650 with 256 GB of RAM, while Canu was run on a cluster with an AMD Epyc 7551P and 256 Gb of RAM. NextDenovo ran in a few hours, while Canu runs each required several days.

## Declarations

### Ethics approval and consent to participate

Not applicable.

### Consent for publication

Not applicable.

### Availability of data and materials

The datasets analyzed during the current study were published in Simion et al. (2020).

### Authors’ contributions

N.G. and J-F.F jointly devised the study. N.G., A.H. and A.D. ran the assemblies. N.G. evaluated the assemblies. A.H. conceived and implemented HapPy. N.G. and J-F.F. wrote the manuscript. K.V.D. and J-F.F. provided the sequencing data. All authors took part in revising the manuscript.

## Acknowledgements

We thank Antoine Limasset and Paul Simion for their useful advice. We also thank Michael Eitel for prompting us to initiate this benchmarking of long-read assemblers. Nanopore reads were generated at Genoscope as part of the France Génomique project ‘ALPAGA’ coordinated by Etienne Danchin (www.france-genomique.org/projet/alpaga/). Part of this analysis was performed on computing clusters of the Leibniz-Rechenzentrum (LRZ) and the Consortium des Équipements de Calcul Intensif (CÉCI) funded by the Fonds de la Recherche Scientifique de Belgique (F.R.S.-FNRS) under Grant No. 2.5020.11.

