## Supplementary_data for "Overcoming uncollapsed haplotypes in long-read assemblies of non-model organisms"

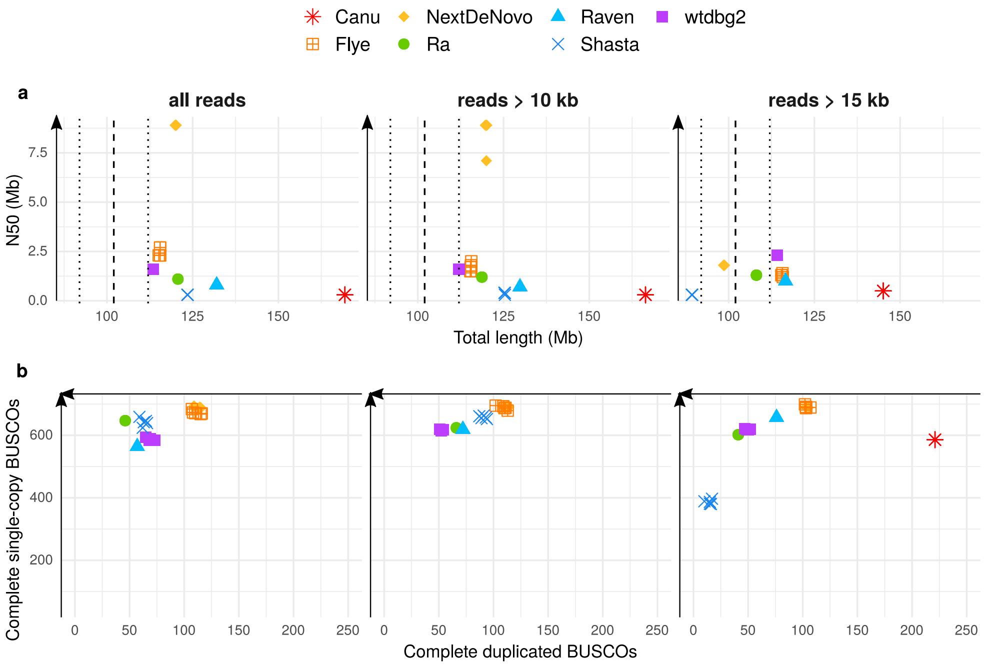


**Figure S1.** Statistics of PacBio assemblies obtained from the full PacBio dataset or with a read-filtering step prior to assembly, using different thresholds: 10 kb, 15 kb. All assemblies were run five times to assess the reproducibility of the output produced by each assembler. A) N50 plotted against total assembly length. B) Number of complete single-copy BUSCOs plotted against number of complete duplicated BUSCOs, from a total of 954 orthologs.

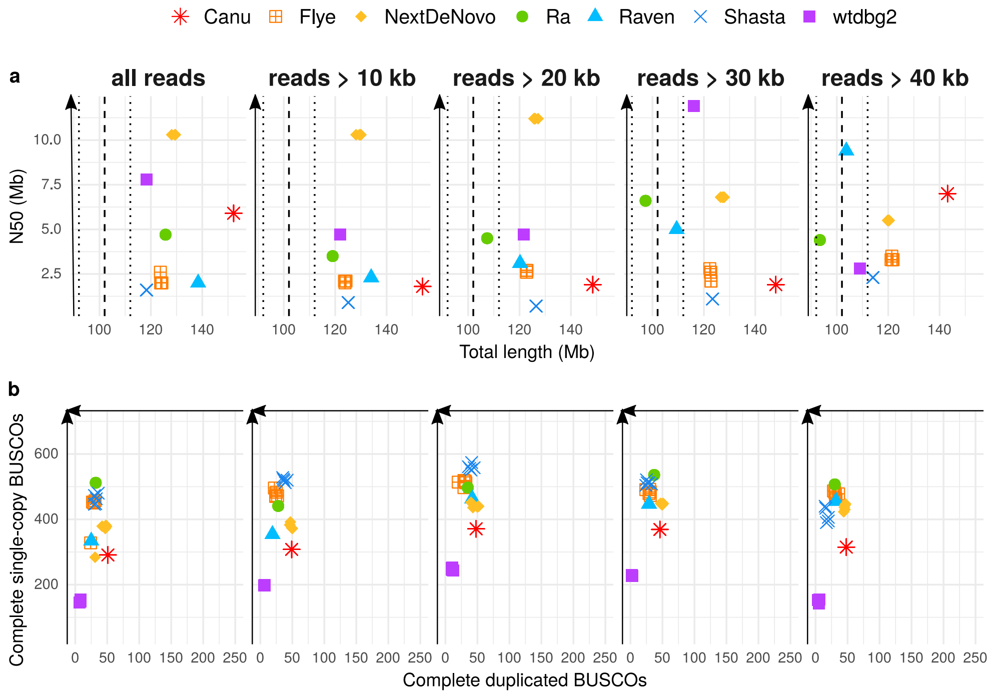


**Figure S2.** Statistics of Nanopore assemblies obtained from the full Nanopore dataset or with a read-filtering step prior to assembly, using different thresholds: 10 kb, 20 kb, 30 kb, 40 kb. All assemblies were run five times to assess the reproducibility of the output produced by each assembler. A) N50 plotted against total assembly length. B) Number of complete single-copy BUSCOs plotted against number of complete duplicated BUSCOs, from a total of 954 orthologs.

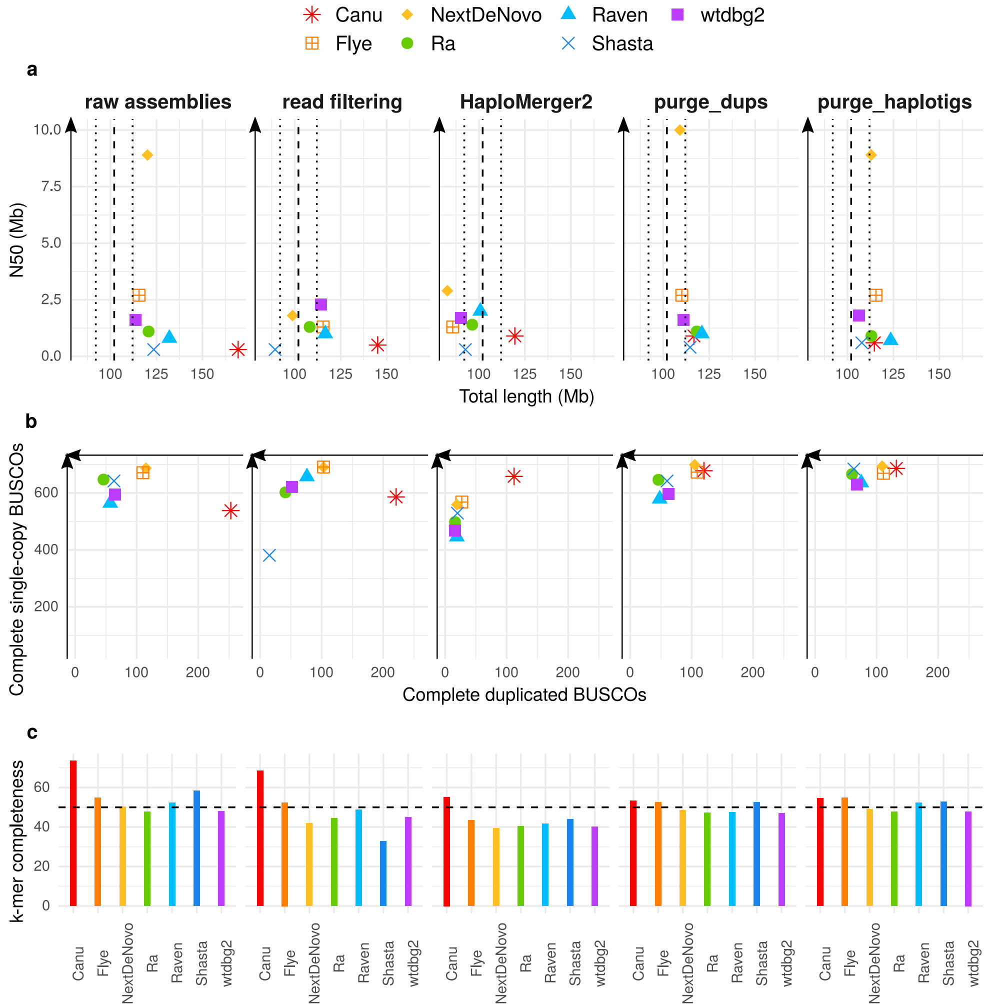


**Figure S3.** Statistics of PacBio assemblies obtained from the full PacBio dataset. a) N50 plotted against total assembly length. b) Number of complete single-copy BUSCOs plotted against number of complete duplicated BUSCOs, from a total of 954 orthologs. c) *k*-mer completeness.

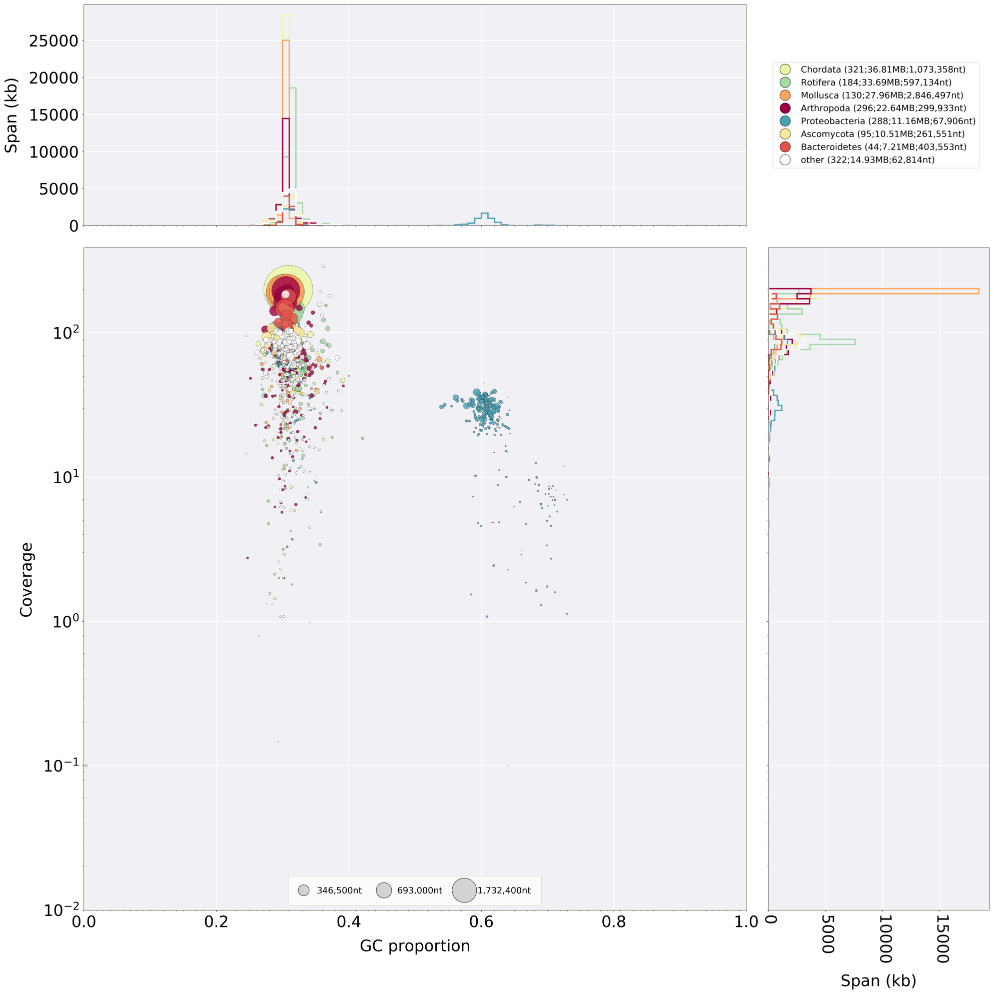


**Figure S4.** Blobtools (Challis et al., 2020) analysis of a Canu assembly of the full PacBio dataset.

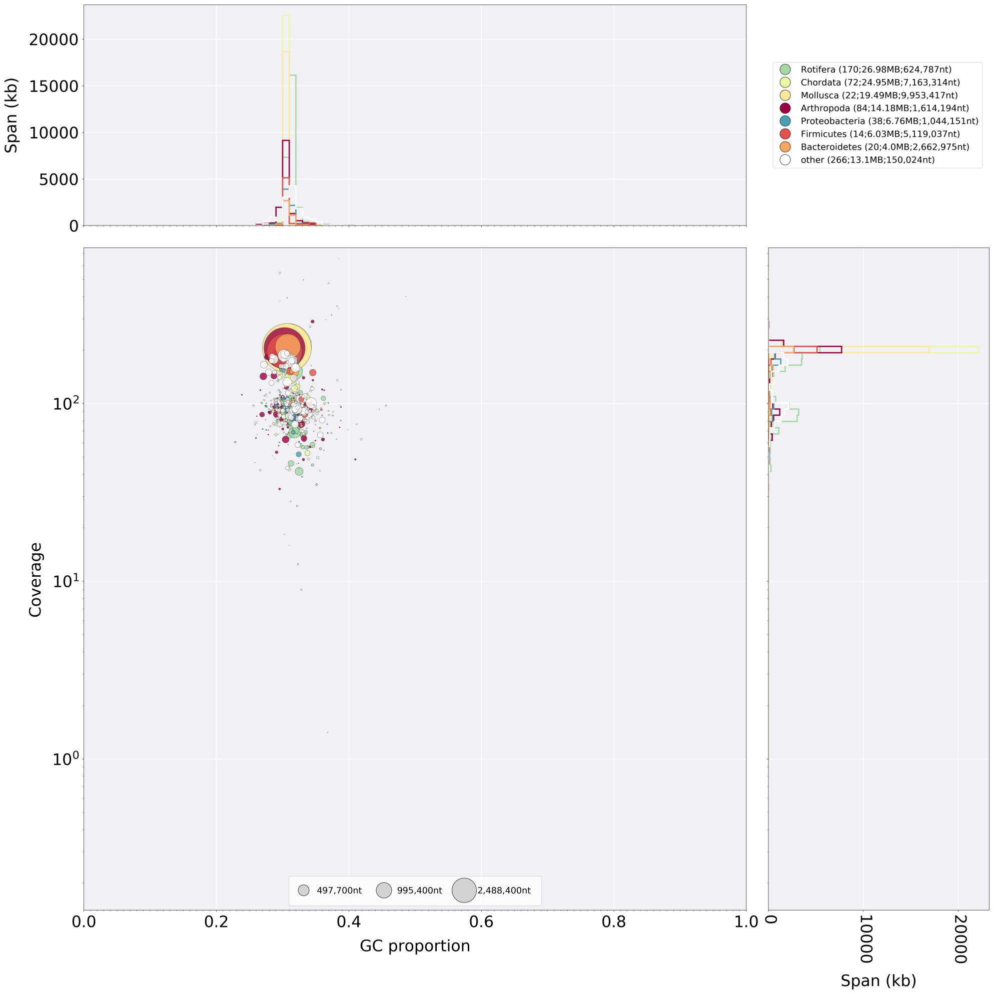


**Figure S5.** Blobtools analysis of a Flye assembly of the full PacBio dataset.

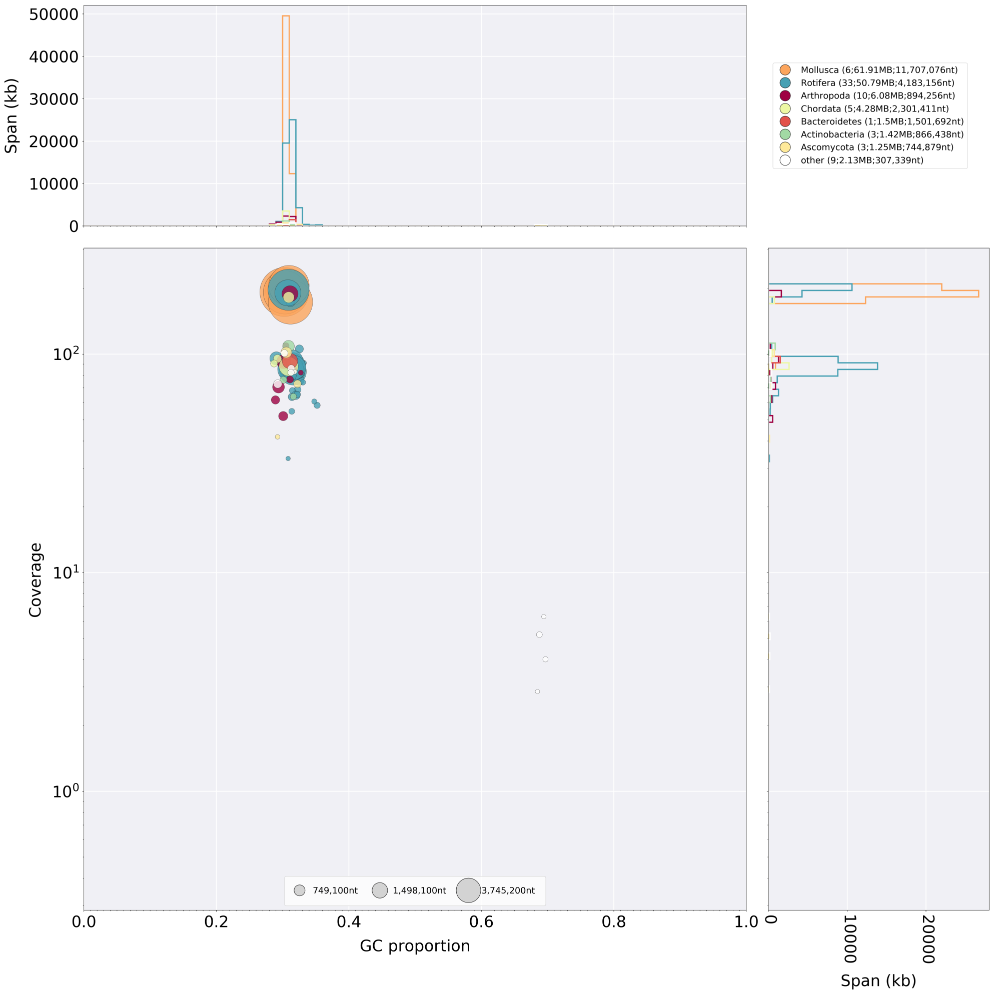


**Figure S6.** Blobtools analysis of a NextDenovo assembly of the full PacBio dataset.

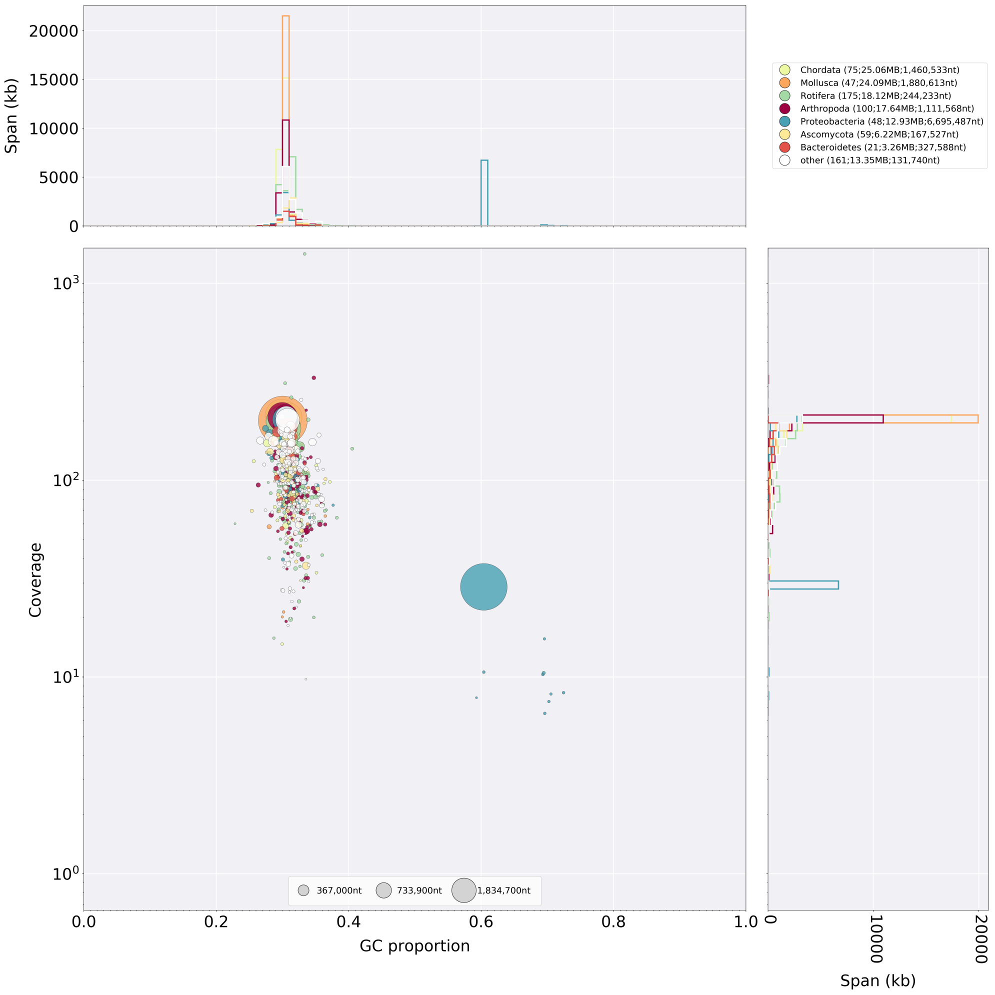


**Figure S7.** Blobtools analysis of a Ra assembly of the full PacBio dataset.

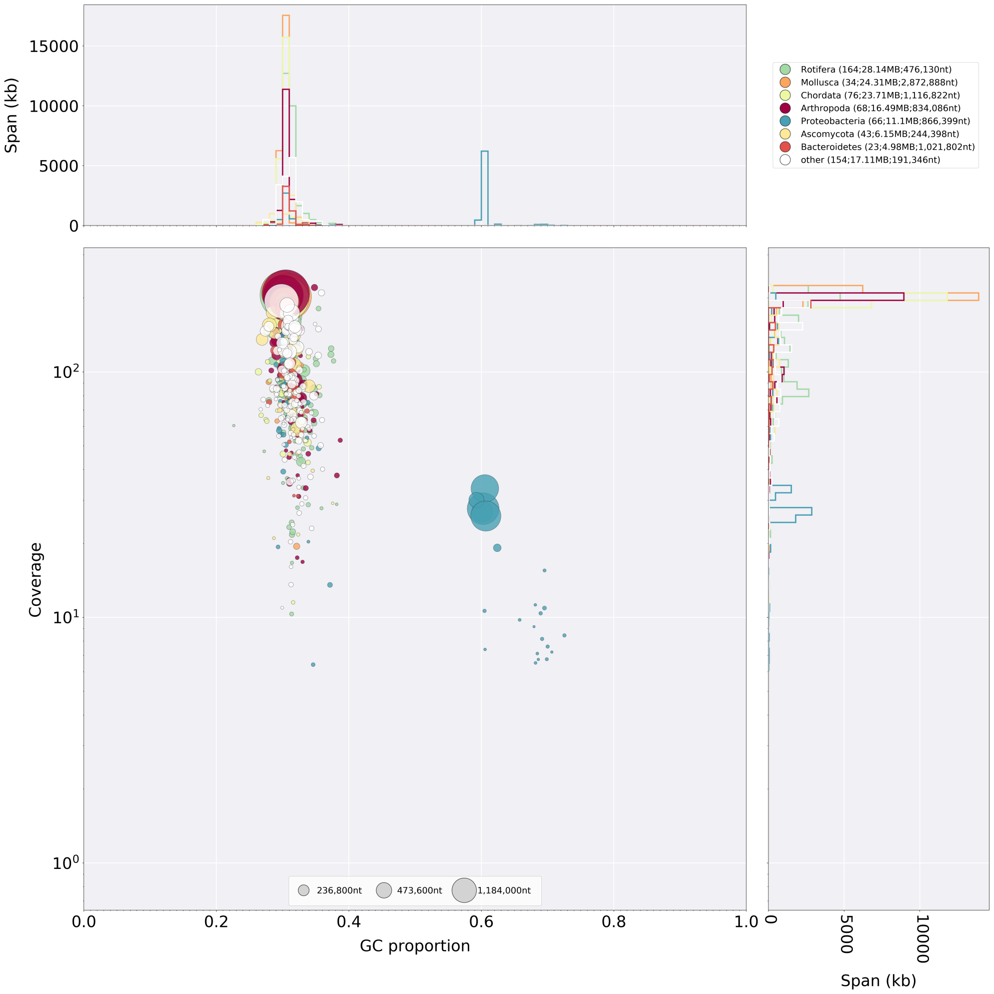


**Figure S8.** Blobtools analysis of a Raven assembly of the full PacBio dataset.

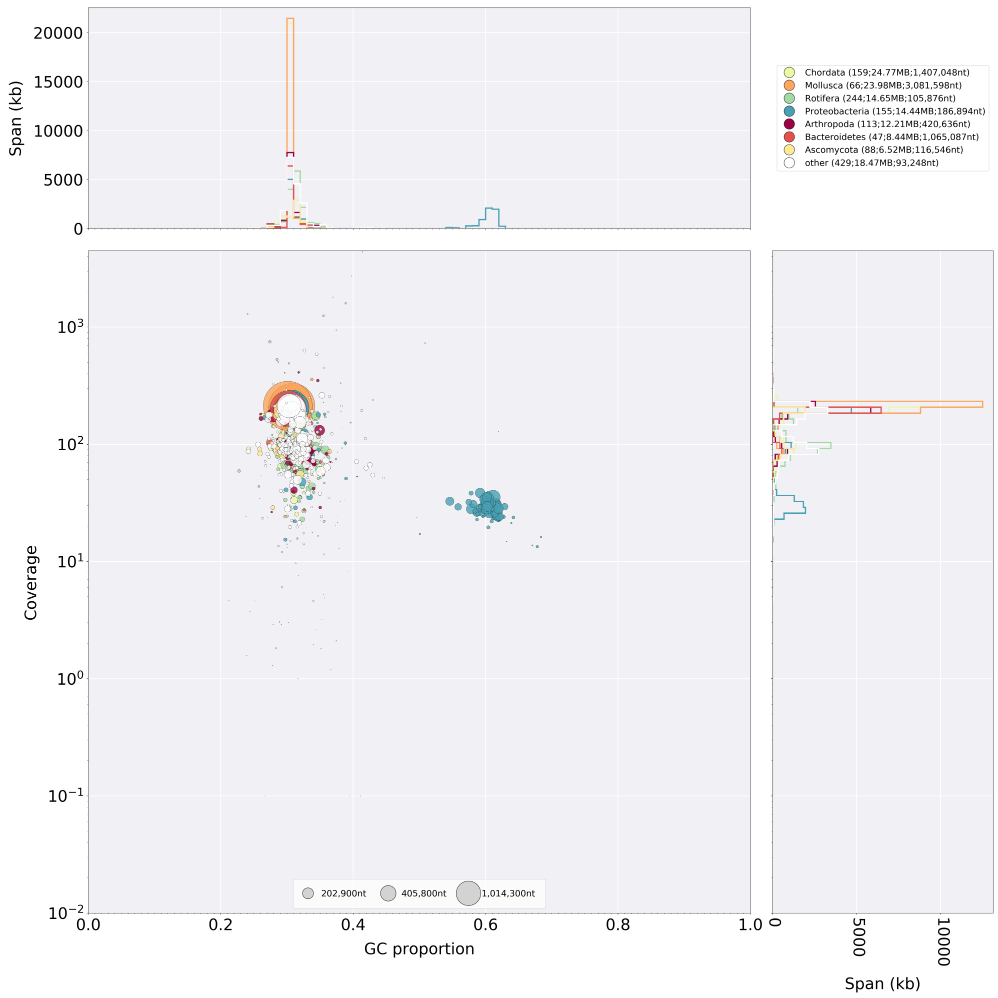


**Figure S9.** Blobtools analysis of a Shasta assembly of the full PacBio dataset.

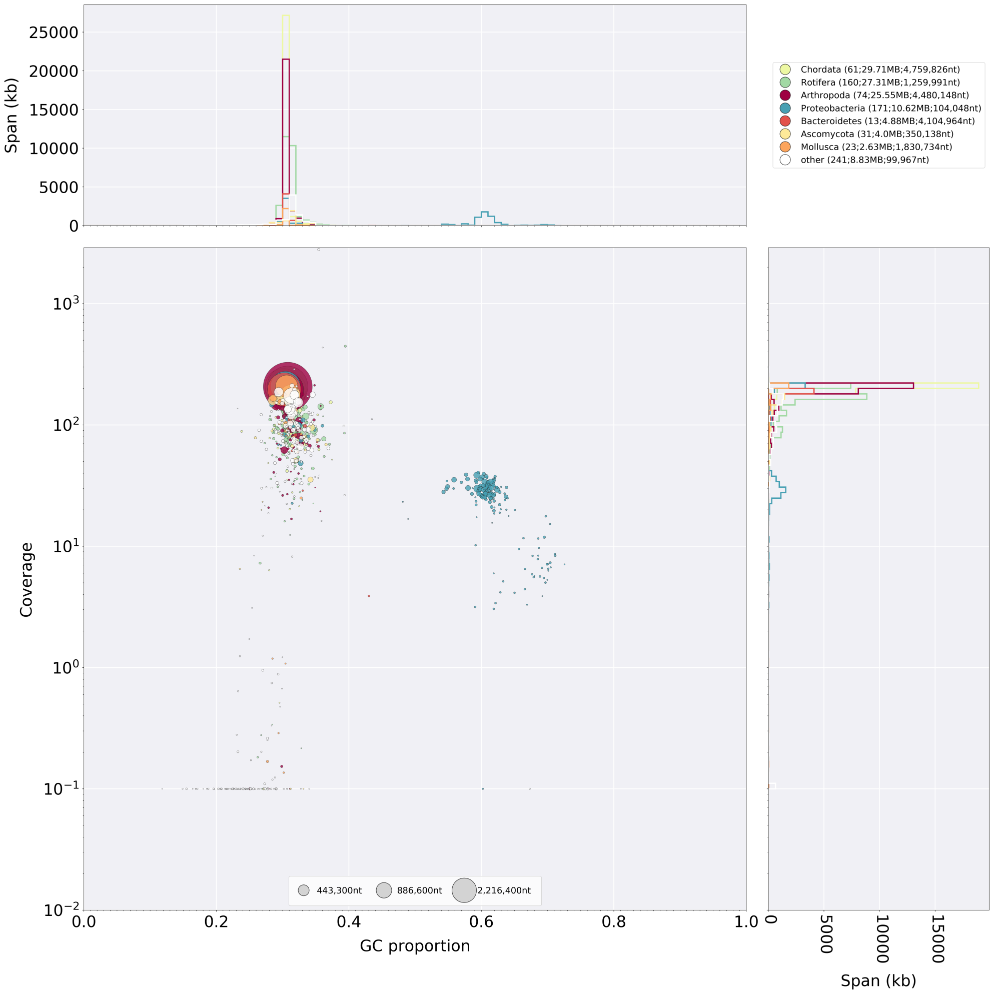


**Figure S10.** Blobtools analysis of a wtdbg2 assembly of the full PacBio dataset.

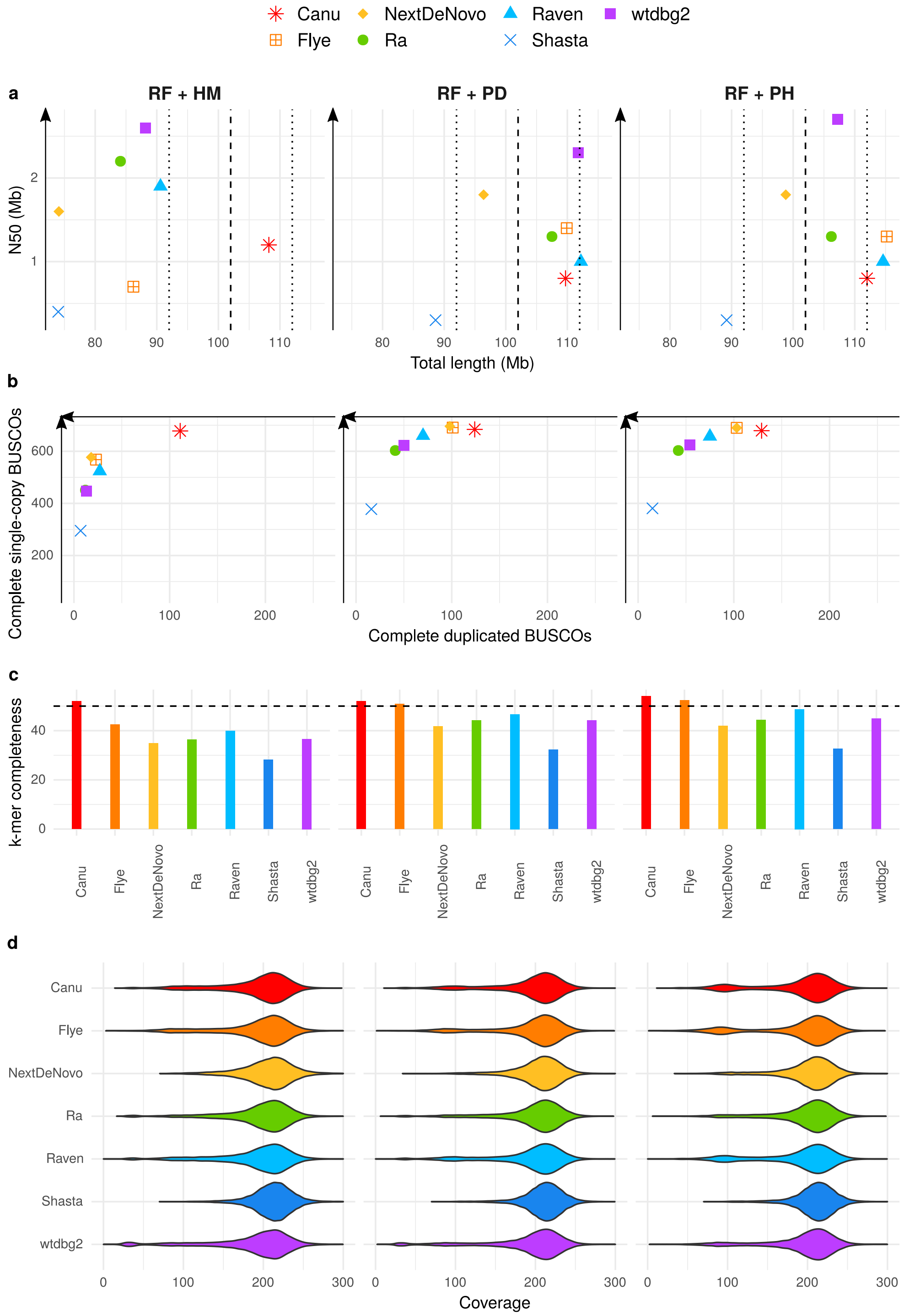


**Figure S11.** Statistics of PacBio assemblies obtained from the filtered PacBio dataset of reads longer than 15 kb, with a subsequent removal of uncollapsed haplotypes with HaploMerger2 (HM), purge_dups (PD), or purge_haplotigs (PH). a) N50 plotted against total assembly length. b) Number of complete single-copy BUSCOs plotted against number of complete duplicated BUSCOs, from a total of 954 orthologs. c) *k*-mer completeness. d) Long-read coverage distribution over the contigs.

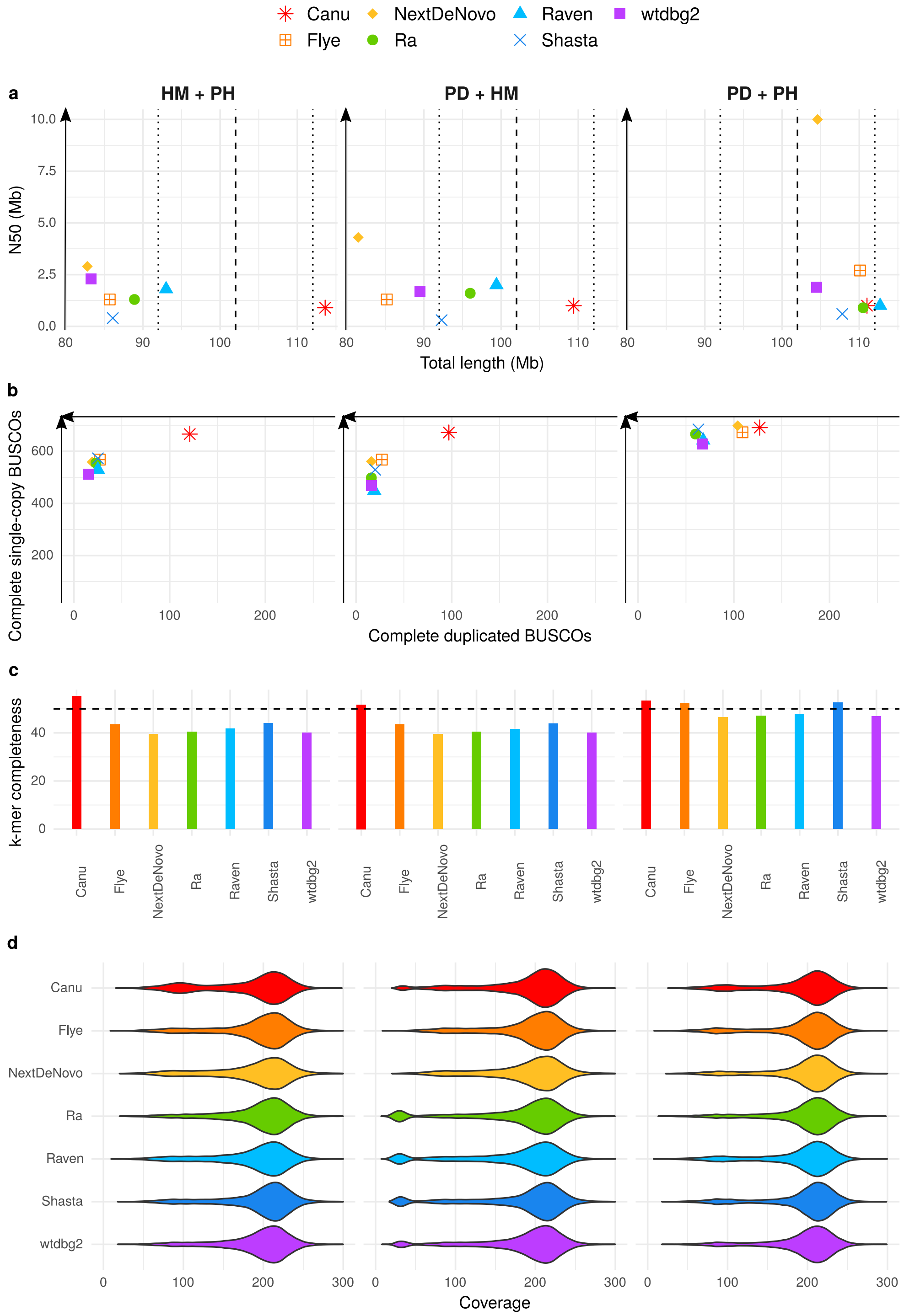


**Figure S12.** Statistics of PacBio assemblies obtained from the full PacBio dataset with a subsequent removal of uncollapsed haplotypes with combinations of HaploMerger2 (HM), purge_dups (PD), and/or purge_haplotigs (PH). a) N50 plotted against total assembly length. b) Number of complete single-copy BUSCOs plotted against number of complete duplicated BUSCOs, from a total of 954 orthologs. c) *k*-mer completeness. d) Long-read coverage distribution over the contigs.

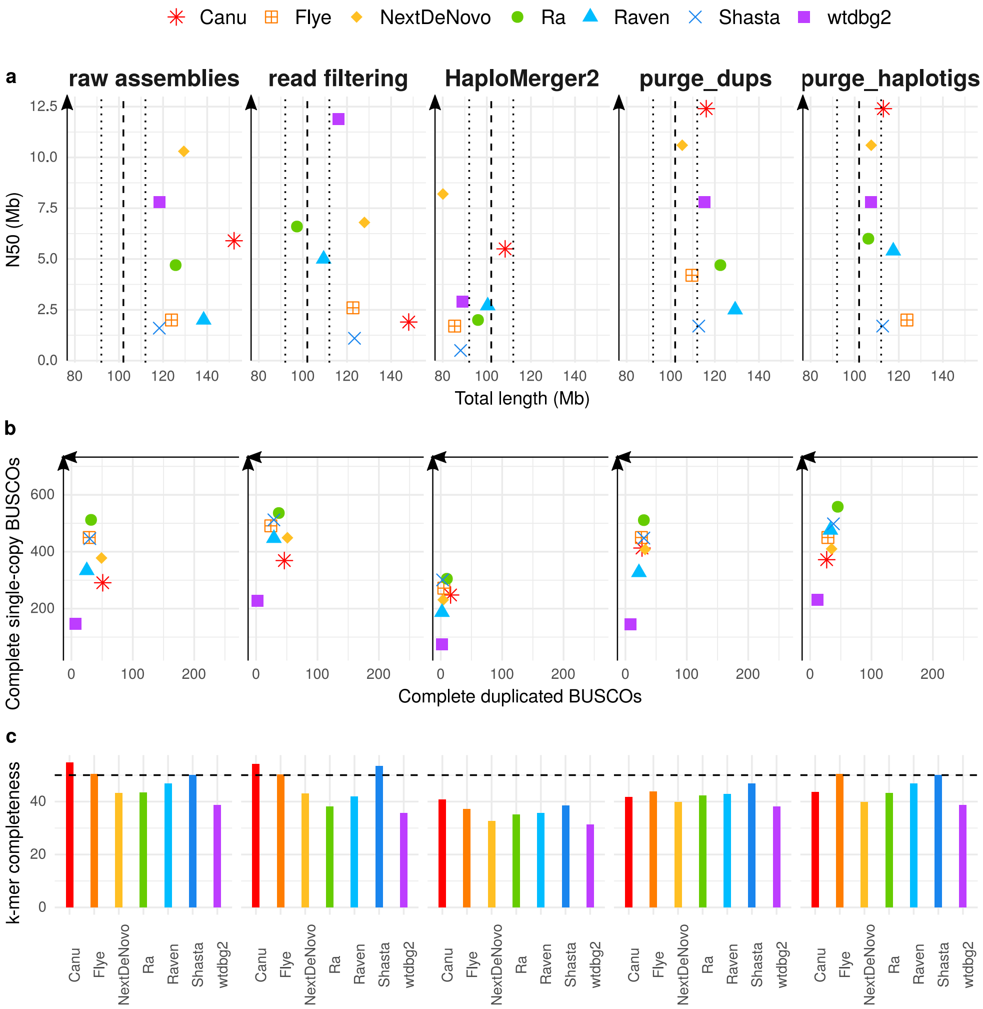


**Figure S13.** Statistics of Nanopore assemblies obtained from the full Nanopore dataset. a) N50 plotted against total assembly length. b) Number of complete single-copy BUSCOs plotted against number of complete duplicated BUSCOs, from a total of 954 orthologs. c) *k*-mer completeness.

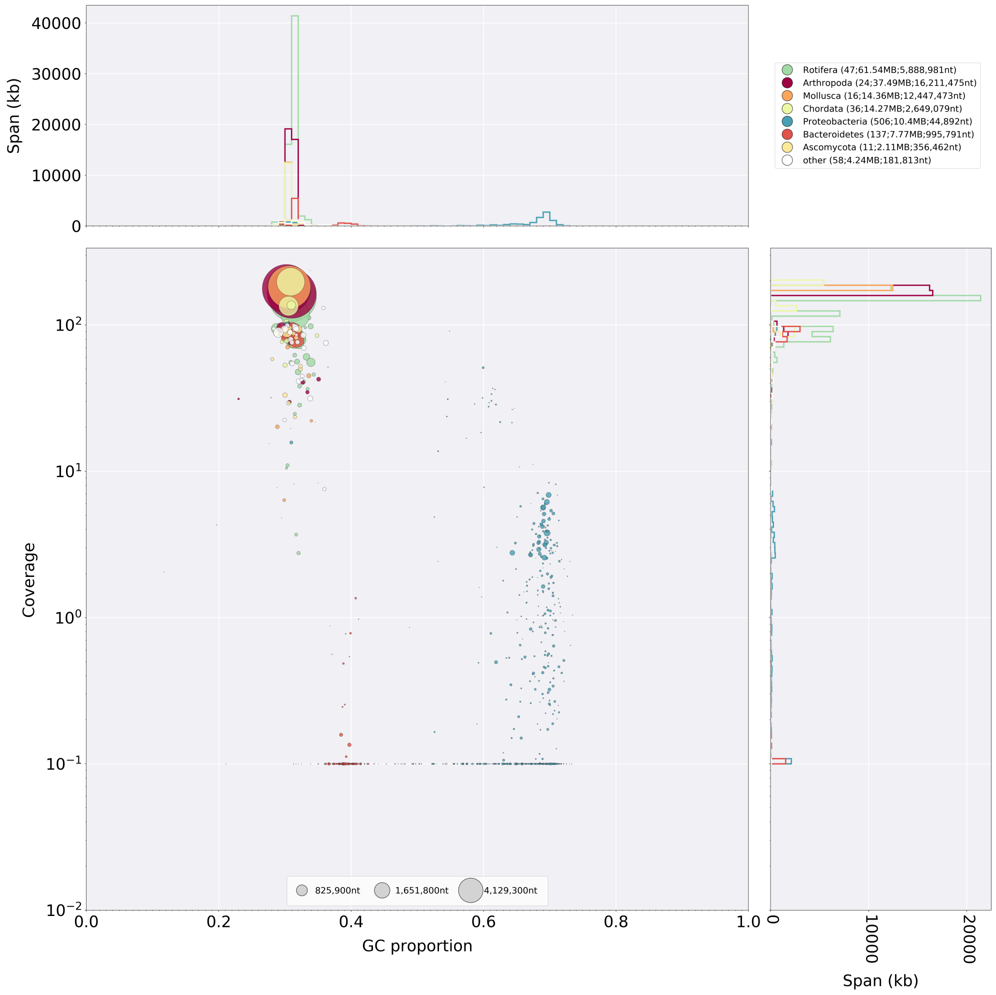


**Figure S14.** Blobtools analysis of a Canu assembly of the full Nanopore dataset.

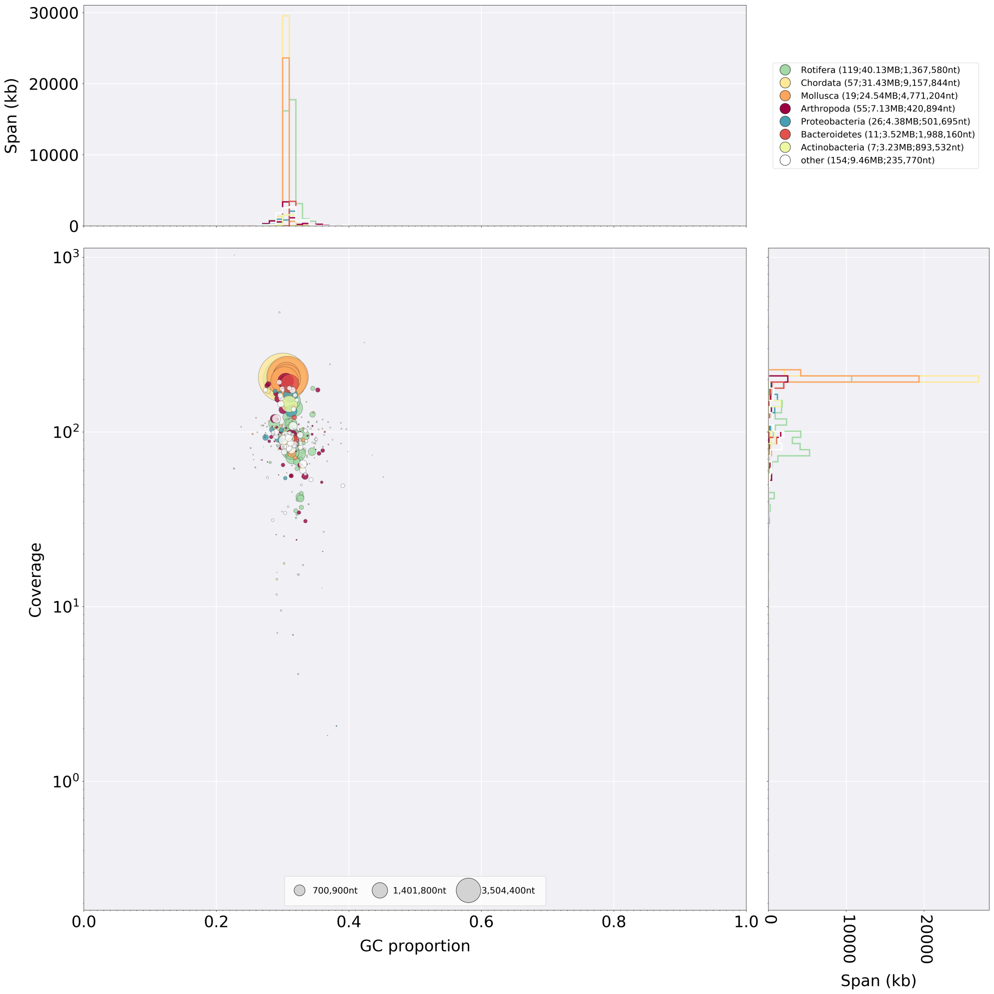


**Figure S15.** Blobtools analysis of a Flye assembly of the full Nanopore dataset.

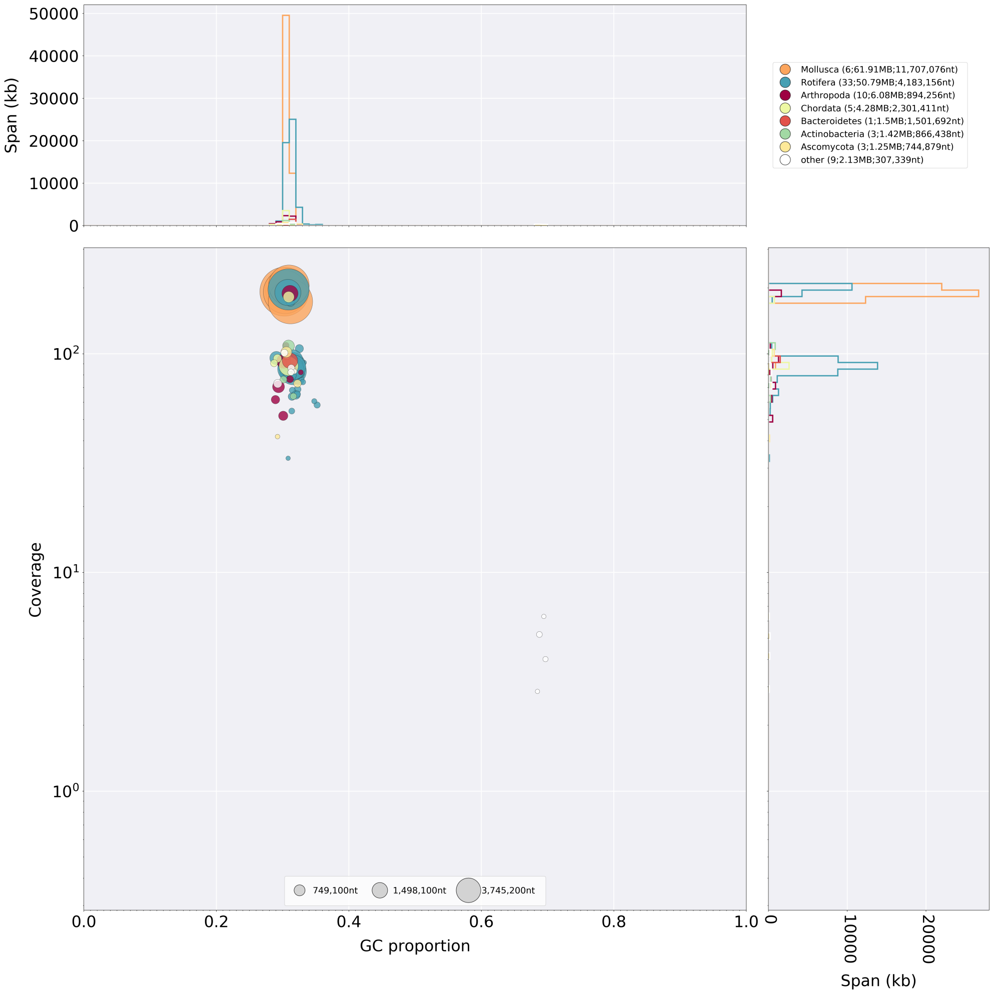


**Figure S16.** Blobtools analysis of a NextDenovo assembly of the full Nanopore dataset.

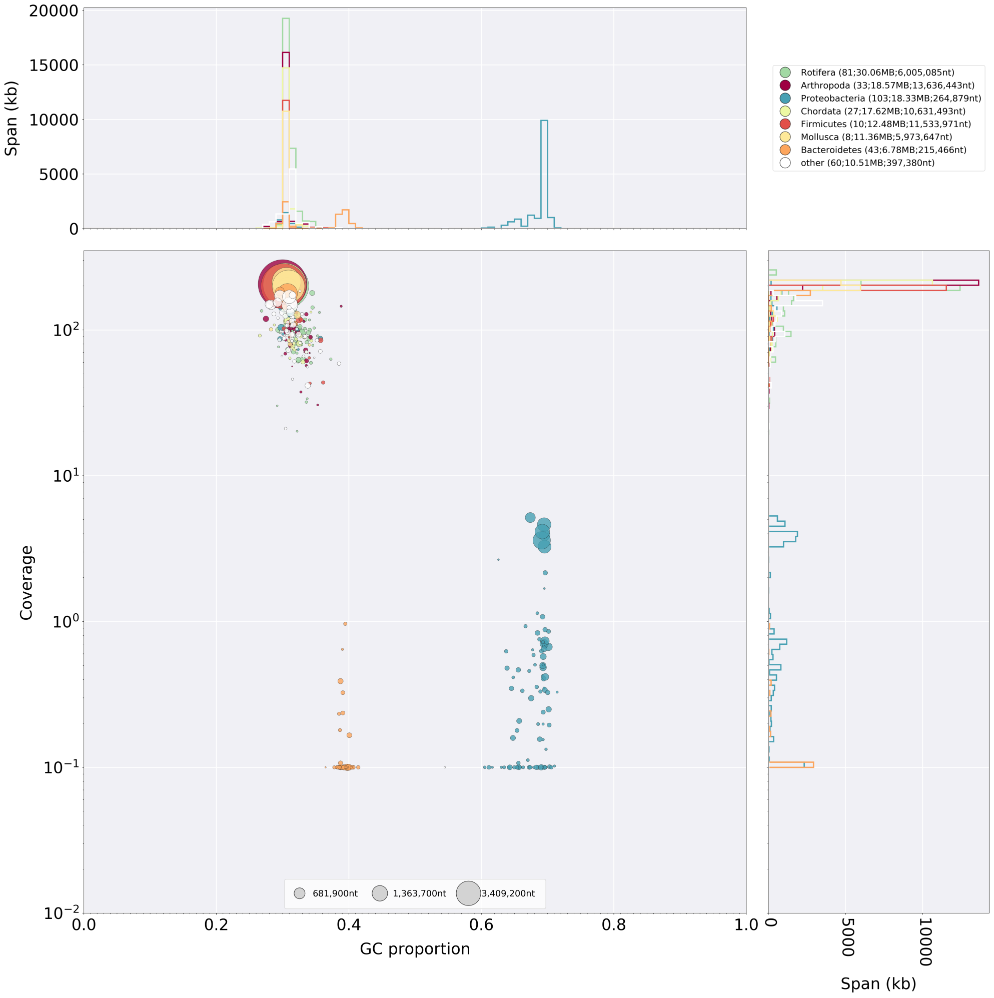


**Figure S17.** Blobtools analysis of a Ra assembly of the full Nanopore dataset.

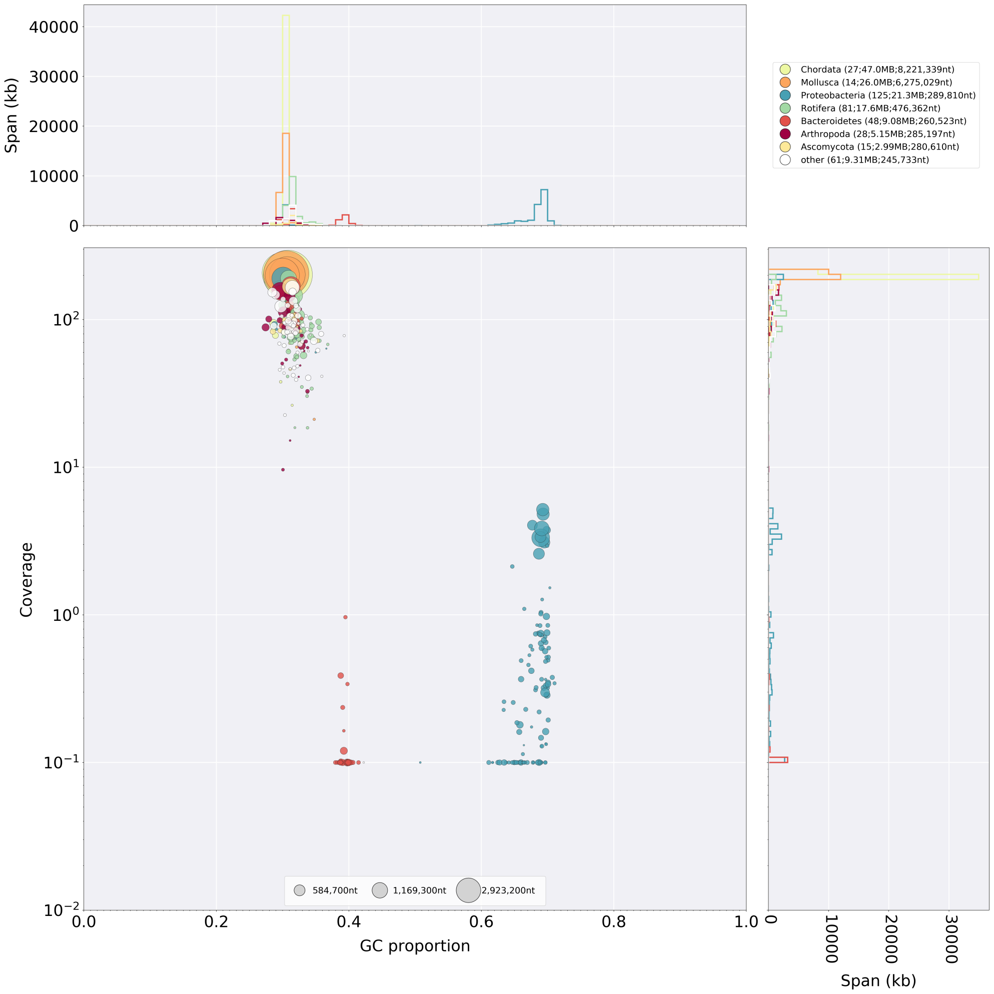


**Figure S18.** Blobtools analysis of a Raven assembly of the full Nanopore dataset.

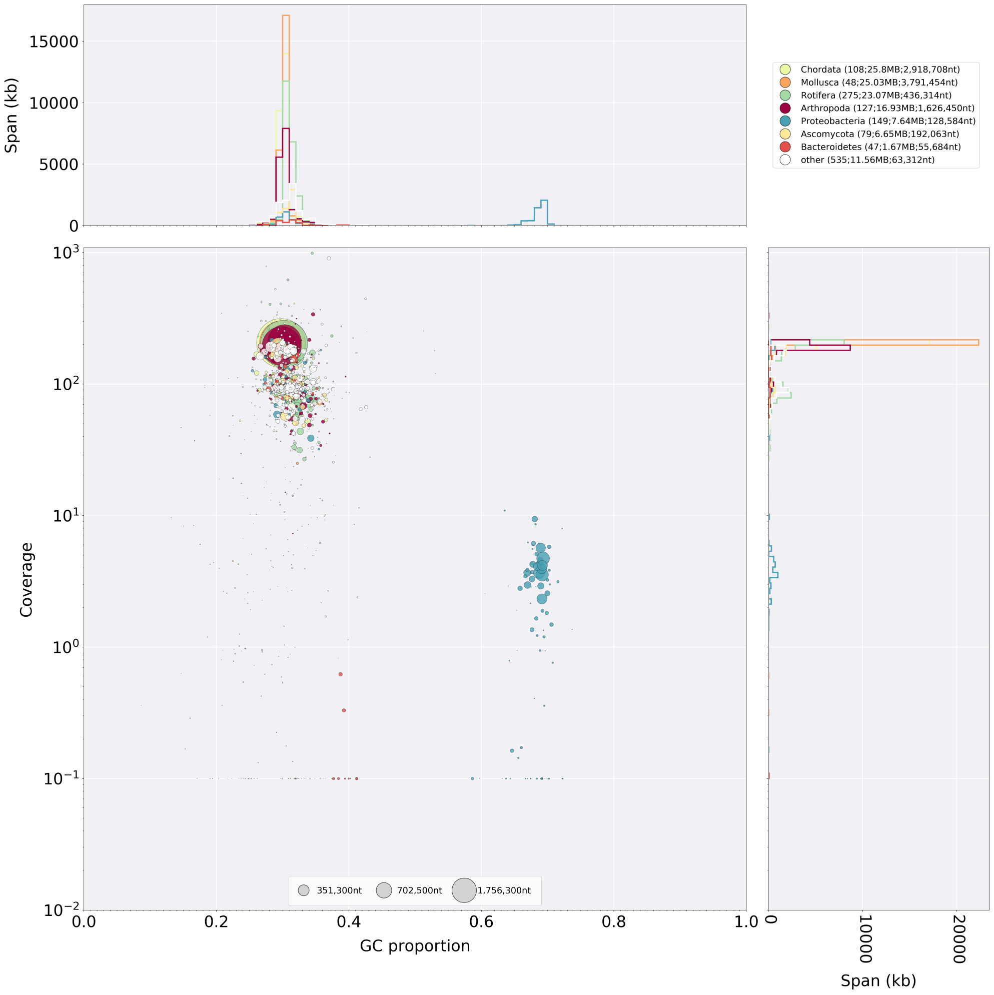


**Figure S19.** Blobtools analysis of a Shasta assembly of the full Nanopore dataset.

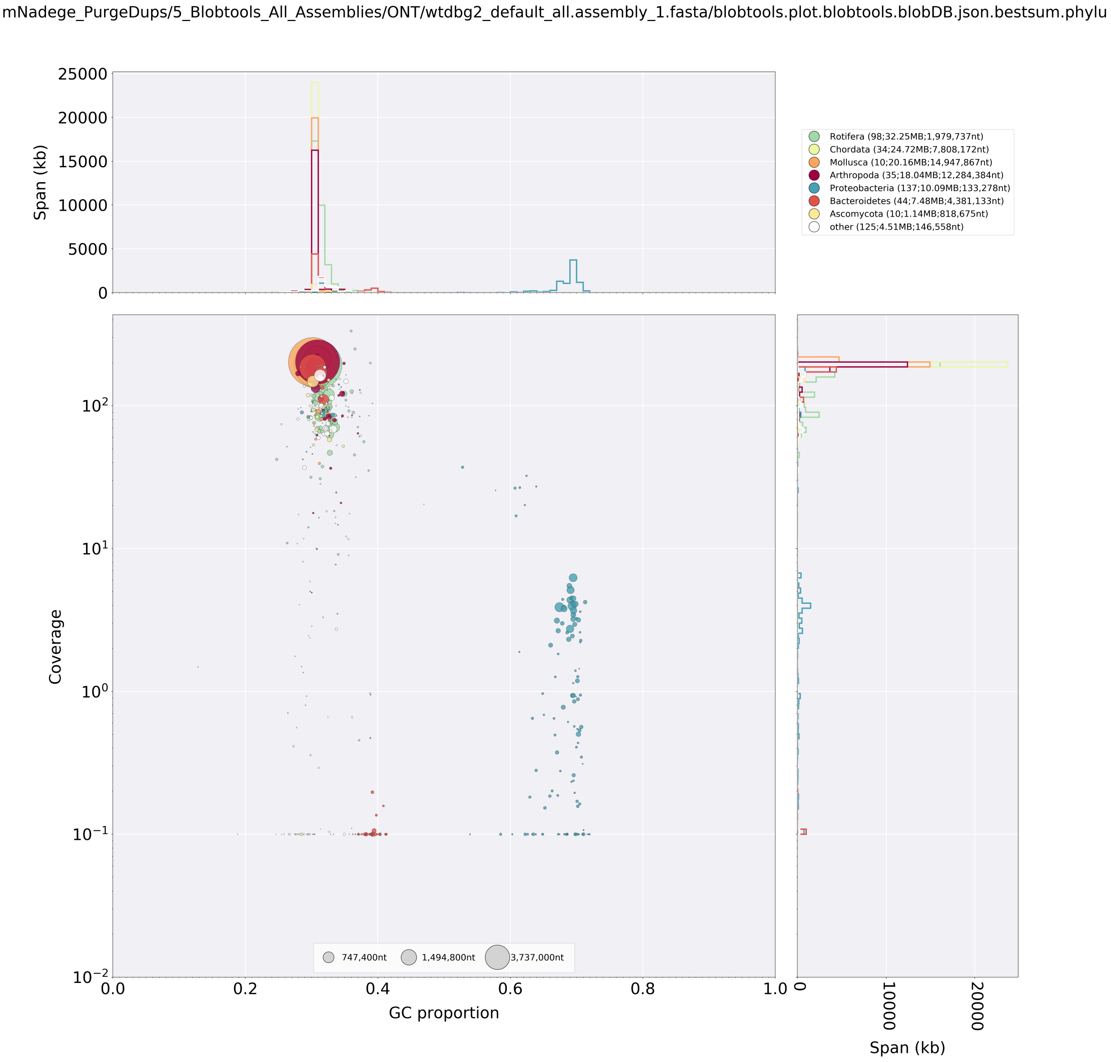


**Figure S20.** Blobtools analysis of a wtdbg2 assembly of the full Nanopore dataset.

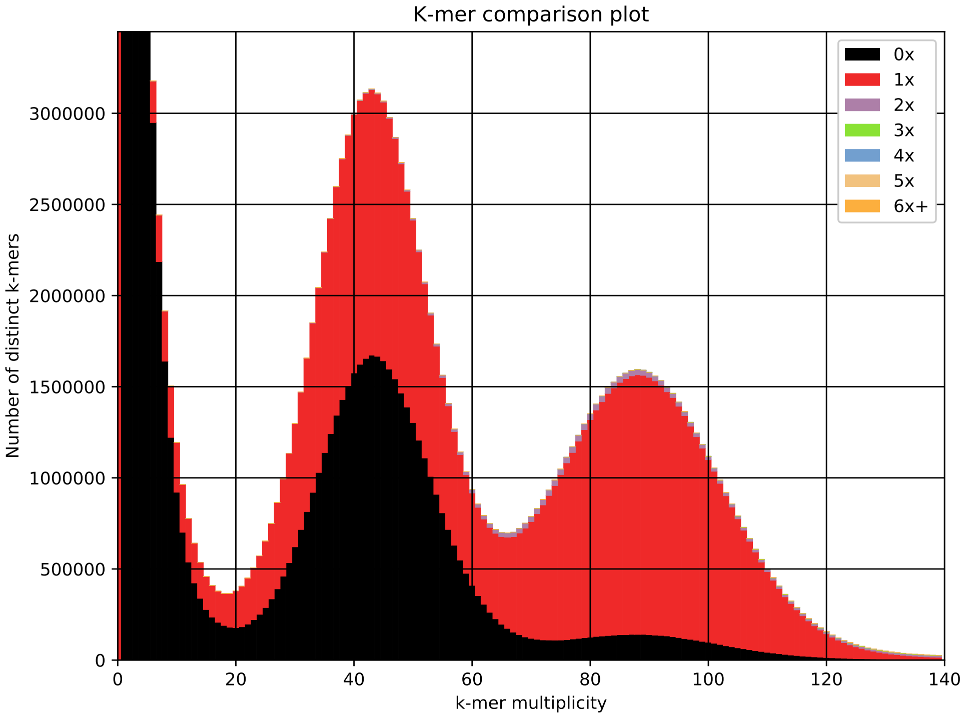


**Figure S21.** *k*-mer spectrum of the Shasta assembly of the full Nanopore dataset.

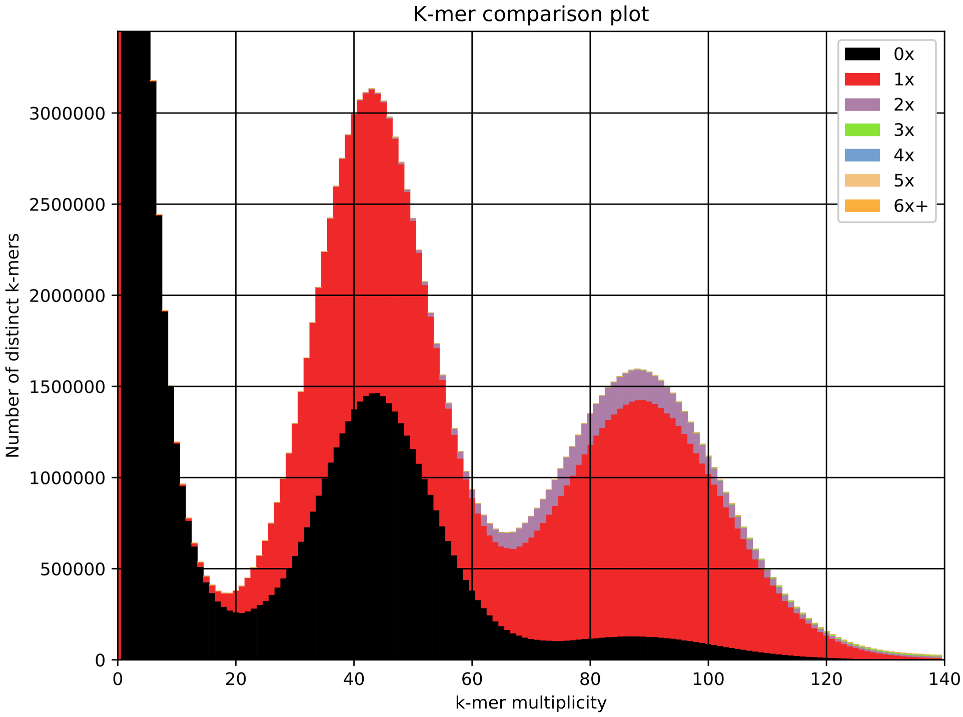


**Figure S22.** *k*-mer spectrum of the Shasta assembly of the longest Nanopore reads.

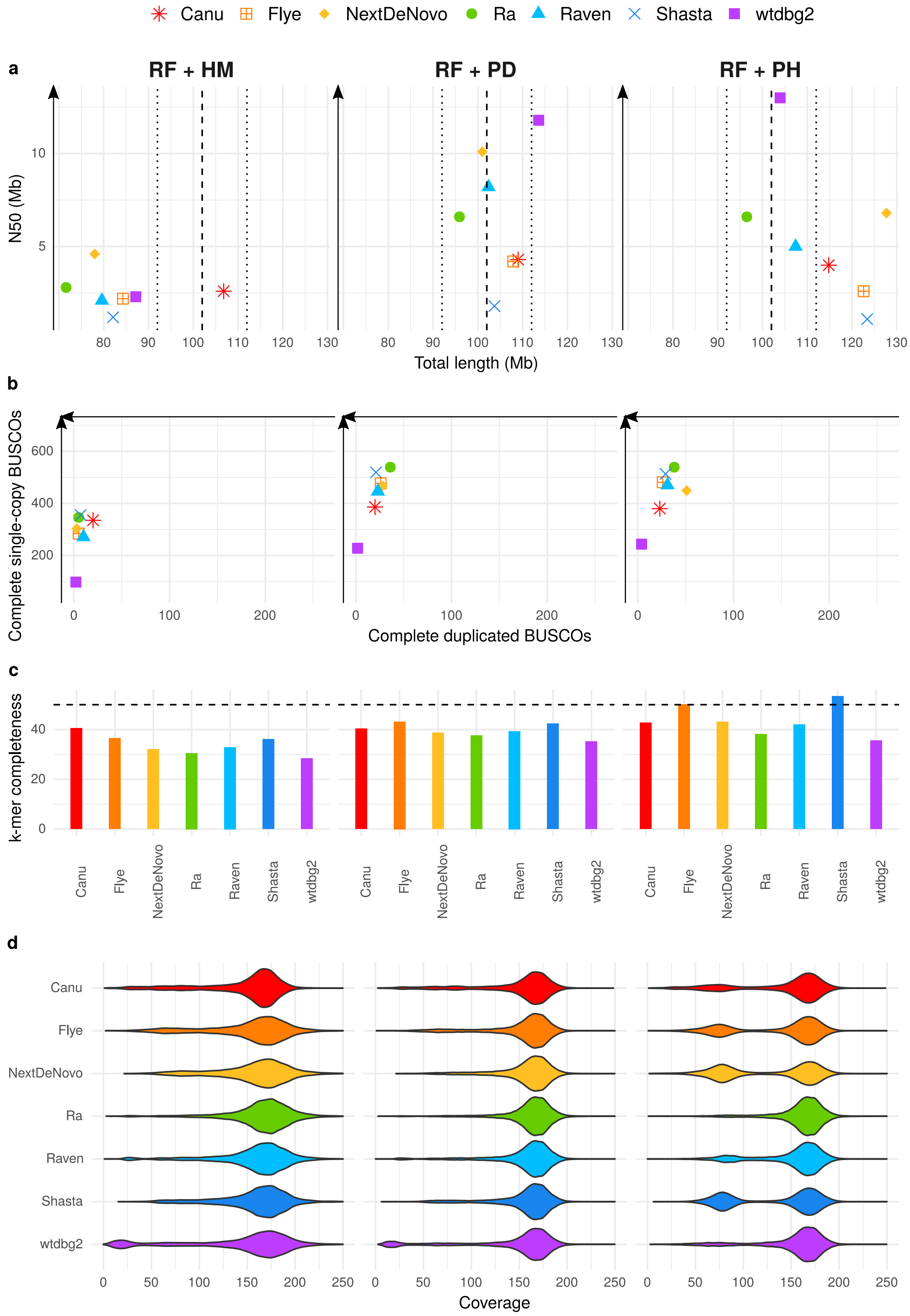


**Figure S23.** Statistics of Nanopore assemblies obtained from the filtered Nanopore dataset of reads longer than 30 kb, with a subsequent removal of uncollapsed haplotypes with HaploMerger2 (HM), purge_dups (PD), or purge_haplotigs (PH). a) N50 plotted against total assembly length. b) Number of complete single-copy BUSCOs plotted against number of complete duplicated BUSCOs, from a total of 954 orthologs. c) *k*-mer completeness. d) Long-read coverage distribution over the contigs.

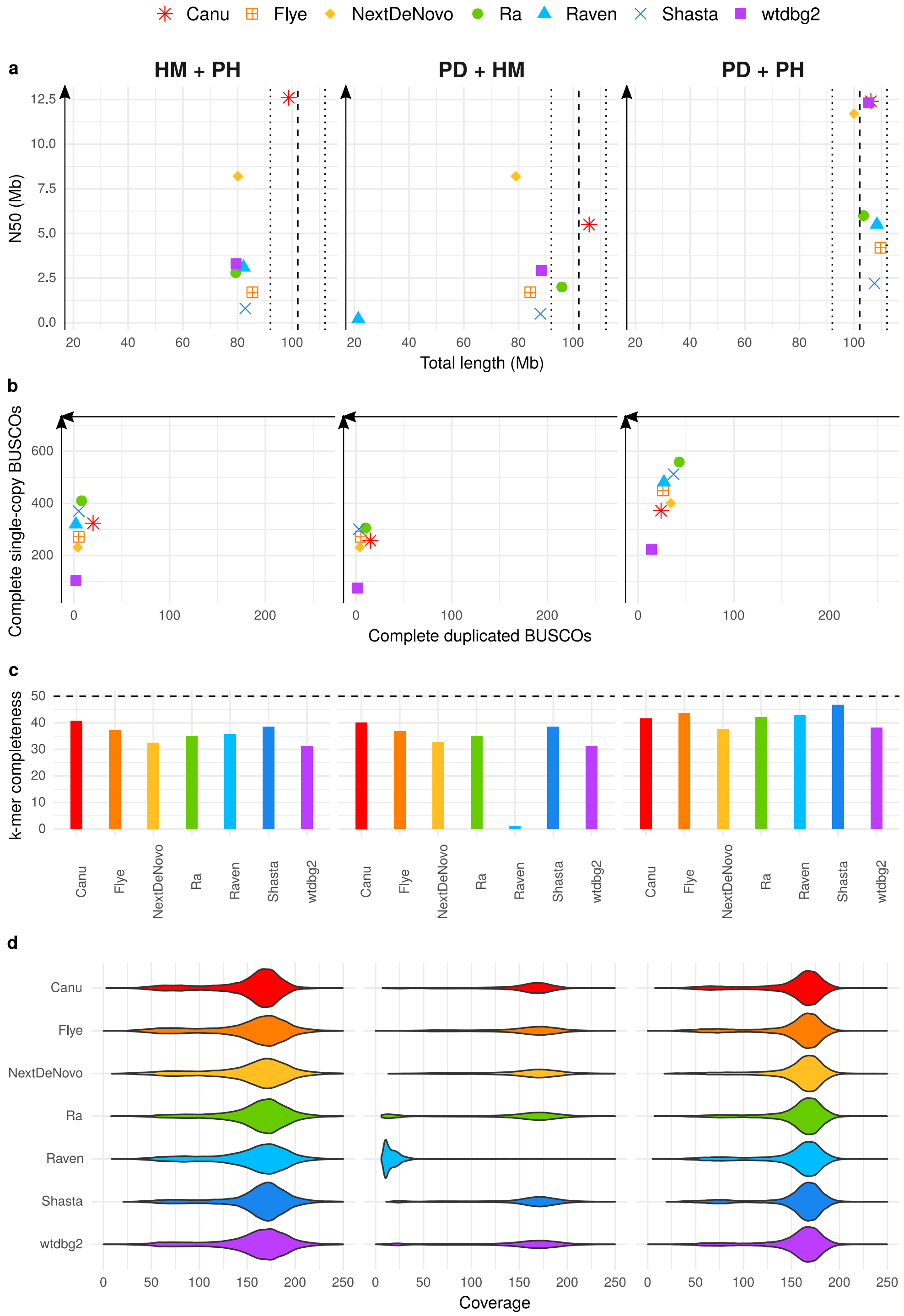


**Figure S24.** Statistics of Nanopore assemblies obtained from the full Nanopore dataset with a subsequent removal of uncollapsed haplotypes with combinations of HaploMerger2 (HM), purge_dups (PD), and/or purge_haplotigs (PH). a) N50 plotted against total assembly length. b) Number of complete single-copy BUSCOs plotted against number of complete duplicated BUSCOs, from a total of 954 orthologs. c) *k*-mer completeness. d) Long-read coverage distribution over the contigs.

**Table S1.** Haploidy values computed by HapPy for PacBio assemblies.

| Assembler | Processing | Haploidy |
|---|---|---|
| Canu | raw assemblies | 0.59 |
| Flye | raw assemblies | 0.85 |
| NextDenovo | raw assemblies | 0.81 |
| Ra | raw assemblies | 0.90 |
| Raven | raw assemblies | 0.82 |
| Shasta | raw assemblies | 0.83 |
| wtdbg2 | raw assemblies | 0.90 |
| Canu | read filtering | 0.62 |
| Flye | read filtering | 0.85 |
| NextDenovo | read filtering | 0.94 |
| Ra | read filtering | 0.94 |
| Raven | read filtering | 0.88 |
| Shasta | read filtering | 0.96 |
| wtdbg2 | read filtering | 0.90 |
| Canu | HaploMerger2 | 0.84 |
| Flye | HaploMerger2 | 0.89 |
| NextDenovo | HaploMerger2 | 0.88 |
| Ra | HaploMerger2 | 0.92 |
| Raven | HaploMerger2 | 0.90 |
| Shasta | HaploMerger2 | 0.91 |
| wtdbg2 | HaploMerger2 | 0.92 |
| Canu | purge_dups | 0.89 |
| Flye | purge_dups | 0.89 |
| NextDenovo | purge_dups | 0.90 |
| Ra | purge_dups | 0.91 |
| Raven | purge_dups | 0.90 |
| Shasta | purge_dups | 0.90 |
| wtdbg2 | purge_dups | 0.91 |
| Canu | purge_haplotigs | 0.86 |
| Flye | purge_haplotigs | 0.85 |
| NextDenovo | purge_haplotigs | 0.87 |
| Ra | purge_haplotigs | 0.88 |
| Raven | purge_haplotigs | 0.80 |
| Shasta | purge_haplotigs | 0.90 |
| wtdbg2 | purge_haplotigs | 0.90 |

**Table S2.** Haploidy values computed by HapPy for PacBio assemblies.

| Assembler | Processing | Haploidy |
|---|---|---|
| Canu | purge_dups + HaploMerger2 | 0.91 |
| Flye | purge_dups + HaploMerger2 | 0.90 |
| NextDenovo | purge_dups + HaploMerger2 | 0.90 |
| Ra | purge_dups + HaploMerger2 | 0.92 |
| Raven | purge_dups + HaploMerger2 | 0.93 |
| Shasta | purge_dups + HaploMerger2 | 0.92 |
| wtdbg2 | purge_dups + HaploMerger2 | 0.92 |
| Canu | read filtering + purge_haplotigs | 0.87 |
| Flye | read filtering + purge_haplotigs | 0.85 |
| NextDenovo | read filtering + purge_haplotigs | 0.94 |
| Ra | read filtering + purge_haplotigs | 0.92 |
| Raven | read filtering + purge_haplotigs | 0.87 |
| Shasta | read filtering + purge_haplotigs | 0.96 |
| wtdbg2 | read filtering + purge_haplotigs | 0.90 |
| Canu | read filtering + purge_dups | 0.91 |
| Flye | read filtering + purge_dups | 0.90 |
| NextDenovo | read filtering + purge_dups | 0.97 |
| Ra | read filtering + purge_dups | 0.95 |
| Raven | read filtering + purge_dups | 0.91 |
| Shasta | read filtering + purge_dups | 0.97 |
| wtdbg2 | read filtering + purge_dups | 0.92 |
| Canu | HaploMerger2 + purge_haplotigs | 0.82 |
| Flye | HaploMerger2 + purge_haplotigs | 0.89 |
| NextDenovo | HaploMerger2 + purge_haplotigs | 0.88 |
| Ra | HaploMerger2 + purge_haplotigs | 0.88 |
| Raven | HaploMerger2 + purge_haplotigs | 0.83 |
| Shasta | HaploMerger2 + purge_haplotigs | 0.88 |
| wtdbg2 | HaploMerger2 + purge_haplotigs | 0.84 |
| Canu | purge_dups + purge_haplotigs | 0.88 |
| Flye | purge_dups + purge_haplotigs | 0.89 |
| NextDenovo | purge_dups + purge_haplotigs | 0.92 |
| Ra | purge_dups + purge_haplotigs | 0.89 |
| Raven | purge_dups + purge_haplotigs | 0.88 |
| Shasta | purge_dups + purge_haplotigs | 0.90 |
| wtdbg2 | purge_dups + purge_haplotigs | 0.91 |

**Table S3.** Haploidy values computed by HapPy for Nanopore assemblies.

| Assembler | Processing | Haploidy |
|---|---|---|
| Canu | raw assemblies | 0.63 |
| Flye | raw assemblies | 0.79 |
| NextDenovo | raw assemblies | 0.72 |
| Ra | raw assemblies | 0.90 |
| Raven | raw assemblies | 0.83 |
| Shasta | raw assemblies | 0.86 |
| wtdbg2 | raw assemblies | 0.92 |
| Canu | read filtering | 0.59 |
| Flye | read filtering | 0.79 |
| NextDenovo | read filtering | 0.72 |
| Ra | read filtering | 0.95 |
| Raven | read filtering | 0.89 |
| Shasta | read filtering | 0.75 |
| wtdbg2 | read filtering | 0.92 |
| Canu | HaploMerger2 | 0.89 |
| Flye | HaploMerger2 | 0.87 |
| NextDenovo | HaploMerger2 | 0.89 |
| Ra | HaploMerger2 | 0.91 |
| Raven | HaploMerger2 | 0.88 |
| Shasta | HaploMerger2 | 0.90 |
| wtdbg2 | HaploMerger2 | 0.89 |
| Canu | purge_dups | 0.92 |
| Flye | purge_dups | 0.90 |
| NextDenovo | purge_dups | 0.92 |
| Ra | purge_dups | 0.93 |
| Raven | purge_dups | 0.90 |
| Shasta | purge_dups | 0.91 |
| wtdbg2 | purge_dups | 0.93 |
| Canu | purge_haplotigs | 0.86 |
| Flye | purge_haplotigs | 0.79 |
| NextDenovo | purge_haplotigs | 0.90 |
| Ra | purge_haplotigs | 0.90 |
| Raven | purge_haplotigs | 0.83 |
| Shasta | purge_haplotigs | 0.86 |
| wtdbg2 | purge_haplotigs | 0.91 |

**Table S4.** Haploidy values computed by HapPy for Nanopore assemblies.

| Assembler | Processing | Haploidy |
|---|---|---|
| Canu | read filtering + purge_haplotigs | 0.85 |
| Flye | read filtering + purge_haplotigs | 0.79 |
| NextDenovo | read filtering + purge_haplotigs | 0.72 |
| Ra | read filtering + purge_haplotigs | 0.95 |
| Raven | read filtering + purge_haplotigs | 0.89 |
| Shasta | read filtering + purge_haplotigs | 0.75 |
| wtdbg2 | read filtering + purge_haplotigs | 0.91 |
| Canu | read filtering + purge_dups | 0.89 |
| Flye | read filtering + purge_dups | 0.91 |
| NextDenovo | read filtering + purge_dups | 0.95 |
| Ra | read filtering + purge_dups | 0.96 |
| Raven | read filtering + purge_dups | 0.95 |
| Shasta | read filtering + purge_dups | 0.93 |
| wtdbg2 | read filtering + purge_dups | 0.92 |
| Canu | HaploMerger2 + purge_haplotigs | 0.89 |
| Flye | HaploMerger2 + purge_haplotigs | 0.87 |
| NextDenovo | HaploMerger2 + purge_haplotigs | 0.89 |
| Ra | HaploMerger2 + purge_haplotigs | 0.91 |
| Raven | HaploMerger2 + purge_haplotigs | 0.92 |
| Shasta | HaploMerger2 + purge_haplotigs | 0.90 |
| wtdbg2 | HaploMerger2 + purge_haplotigs | 0.90 |
| Canu | purge_dups + purge_haplotigs | 0.91 |
| Flye | purge_dups + purge_haplotigs | 0.90 |
| NextDenovo | purge_dups + purge_haplotigs | 0.94 |
| Ra | purge_dups + purge_haplotigs | 0.93 |
| Raven | purge_dups + purge_haplotigs | 0.90 |
| Shasta | purge_dups + purge_haplotigs | 0.91 |
| wtdbg2 | purge_dups + purge_haplotigs | 0.92 |
| Canu | purge_dups + HaploMerger2 | 0.90 |
| Flye | purge_dups + HaploMerger2 | 0.88 |
| NextDenovo | purge_dups + HaploMerger2 | 0.90 |
| Ra | purge_dups + HaploMerger2 | 0.91 |
| Raven | purge_dups + HaploMerger2 | 0.51 |
| Shasta | purge_dups + HaploMerger2 | 0.90 |
| wtdbg2 | purge_dups + HaploMerger2 | 0.89 |

**Table S5.** List of command lines used for each tool. Values L, M, H for `purge_haplotigs cov` were selected for each assembly according to the histogram produced by `purge_haplotigs hist`.

<table>
<tr><th>Program</th><th>Dataset</th><th>Command lines</th></tr>
<tr><td>Canu</td><td>PacBio</td><td>canu -d out -p out genomeSize=100m useGrid=false -pacbio-raw pb_data</td></tr>
<tr><td>Canu</td><td>Nanopore</td><td>canu -d out -p out genomeSize=100m useGrid=false -nanopore-raw ont_data</td></tr>
<tr><td>Flye</td><td>PacBio</td><td>flye -o out -g 100m --pacbio-raw pb_data</td></tr>
<tr><td>Flye</td><td>Nanopore</td><td>flye -o out -g 100m --nano-raw ont_data</td></tr>
<tr><td>NextDenovo</td><td>PacBio</td><td>echo pb_data > input.fofn<br>seq_stat input.fofn -g 100Mb -d 150 > stats.txt<br>NextDenovo run.cfg</td></tr>
<tr><td>NextDenovo</td><td>Nanopore</td><td>echo ont_data > input.fofn<br>seq_stat input.fofn -g 100Mb -d 150 > stats.txt<br>NextDenovo run.cfg</td></tr>
<tr><td>Ra</td><td>PacBio</td><td>ra -x pb pb_data > assembly.fasta</td></tr>
<tr><td>Ra</td><td>Nanopore</td><td>ra -x ont ont_data > assembly.fasta</td></tr>
<tr><td>Raven</td><td>-</td><td>raven long_read_data > assembly.fasta</td></tr>
<tr><td>Shasta</td><td>PacBio</td><td>shasta --input pb_data --Reads.minReadLength 0 --assemblyDirectory out --Assembly.consensusCaller Modal --Kmers.k 12</td></tr>
<tr><td>Shasta</td><td>Nanopore</td><td>shasta --input ont_data --Reads.minReadLength 0 --assemblyDirectory out</td></tr>
<tr><td>wtdbg2</td><td>PacBio</td><td>wtdbg2 -x rs -g 100m -i pb_data -fo out<br>wtpoa-cns -i out.ctg.lay.gz -o out.ctg.fa<br>minimap2 -x map-pb -a out.ctg.fa pb_data | samtools sort > out.ctg.bam<br>samtools view out.ctg.bam | wtpoa-cns -d out.ctg.fa -i - -fo assembly.fasta</td></tr>
<tr><td>wtdbg2</td><td>Nanopore</td><td>wtdbg2 -x ont -g 100m -i ont_data -fo out<br>wtpoa-cns -i out.ctg.lay.gz -o out.ctg.fa<br>minimap2 -x map-ont -a out.ctg.fa ont_data | samtools sort > out.ctg.bam<br>samtools view out.ctg.bam | wtpoa-cns -d out.ctg.fa -i - -fo assembly.fasta</td></tr>
<tr><td>HaploMerger2</td><td>-</td><td>samtools faidx assembly.fasta<br>BuildDatabase -name asm.db -engine ncbi assembly.fasta<br>RepeatModeler -engine ncbi -database asm.db<br>RepeatMasker -e ncbi -lib consensi.fa -xsmall assembly.fasta<br>run_all.batch</td></tr>
<tr><td>purge_dups</td><td>PacBio</td><td>echo pb_data > input.fofn<br>pd_config.py assembly.fasta input.fofn<br>run_purge_dups.py config.json purge_dups_bin species_id</td></tr>
<tr><td>purge_dups</td><td>Nanopore</td><td>echo ont_data > input.fofn<br>pd_config.py assembly.fasta input.fofn<br>run_purge_dups.py config.json purge_dups_bin species_id</td></tr>
<tr><td>purge_haplotigs</td><td>PacBio</td><td>minimap2 -ax map-pb assembly.fasta ont_data --secondary=no > aligned.bam<br>samtools sort -o ali.sorted.bam -T tmp.ali aligned.bam<br>samtools index ali.sorted.bam<br>samtools faidx assembly.fasta<br>purge_haplotigs hist -b ali.sorted.bam -g assembly.fasta<br>purge_haplotigs cov -i ali.sorted.bam -l L -m M -h H -o cov_stats.csv<br>purge_haplotigs purge -g assembly.fasta -c cov_stats.csv -o assembly.purged.fasta</td></tr>
<tr><td>purge_haplotigs</td><td>Nanopore</td><td>minimap2 -ax map-ont assembly.fasta ont_data --secondary=no > aligned.bam<br>samtools sort -o ali.sorted.bam -T tmp.ali aligned.bam<br>samtools index ali.sorted.bam<br>samtools faidx assembly.fasta<br>purge_haplotigs hist -b ali.sorted.bam -g assembly.fasta<br>purge_haplotigs cov -i ali.sorted.bam -l L -m M -h H -o cov_stats.csv<br>purge_haplotigs purge -g assembly.fasta -c cov_stats.csv -o assembly.purged.fasta</td></tr>
<tr><td>BBtools</td><td>-</td><td>reformat.sh in=long_reads_data out=subset_data samplebasestarget=number_of_bases</td></tr>
<tr><td>BUSCO</td><td>-</td><td>busco -i assembly.fasta -o busco_output -l metazoa_odb10 -m genome</td></tr>
<tr><td>KAT</td><td>Illumina</td><td>kat comp -o kat_output ’end1.fastq end2.fastq’ assembly.fasta</td></tr>
<tr><td>tinycov</td><td>Nanopore</td><td>minimap2 -x map-ont -a assembly.fasta ont_data | samtools sort > aligned.bam<br>tinycov covplot -r 20000 -t cov.txt aligned.bam</td></tr>
<tr><td>tinycov</td><td>PacBio</td><td>minimap2 -x map-pb -a assembly.fasta pb_data | samtools sort > aligned.bam<br>tinycov covplot -r 20000 -t cov.txt aligned.bam</td></tr>
<tr><td>HapPy</td><td>Nanopore</td><td>minimap2 -x map-ont -a assembly.fasta ont_data | samtools sort > aligned.bam<br>HapPy.py depth aligned.bam out_dir<br>HapPy.py estimate out_dir/aligned.bam.hist</td></tr>
<tr><td>HapPy</td><td>PacBio</td><td>minimap2 -x map-pb -a assembly.fasta pb_data | samtools sort > aligned.bam<br>HapPy.py depth aligned.bam out_dir<br>HapPy.py estimate out_dir/aligned.bam.hist</td></tr>
<tr><td>time</td><td>-</td><td>/usr/bin/time -v -o time_output.txt</td></tr>
</table>

**Table S6.** Long-read and short-read datasets used in the study.

| Data type | Minimum length | Total data | N50 |
|---|---|---|---|
| PacBio | - | 23.5 Gb | 11.6 kb |
| | 15 kb | 4.7 Gb | 17.6 kb |
| Nanopore | - | 17.5 Gb | 18.8 kb |
| | 30 kb | 5.7 Gb | 51.8 kb |
| Illumina 2*250 bp | 30 bp | 11.4 Gb | 250 bp |
